# A generalized test of genotype-phenotype causality in population-sampled nuclear families

**DOI:** 10.64898/2025.12.29.696865

**Authors:** Yushi Tang, John D. Storey

**Affiliations:** Lewis-Sigler Institute for Integrative Genomics, Princeton University, NJ 08544, USA

**Keywords:** causal inference, potential outcomes, nuclear families, randomized experiments, transmission disequilibrium test, transmission mean test

## Abstract

We recently developed a causal inference framework and test — the Transmission Mean Test (TMT) — to identify causal genotype-phenotype relationships in population-sampled parent-child trios, where one child per family is observed. Here, we establish the generalized TMT (gTMT) for population-sampled nuclear families, allowing multiple offspring per family. This extension focuses on detecting genetic loci with non-zero average causal effects (ACE) on child phenotypes, taking into account that siblings share similar random family-specific effects. We construct a potential outcomes trait model that considers both individual-level and family-level heterogeneity, captures additive and non-additive genetic effects, and accommodates both quantitative (continuous or count) and dichotomous traits. We design an unbiased estimate *d*_gTMT_ of the ACE and develop a sampling variance estimate 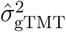 to form a statistic testing the null hypothesis of no causal effect. We provide both theory and empirical evidence demonstrating that gTMT is robust to confounding factors such as the population structure and family-specific effects. We analyze nuclear families in the UK Biobank as an illustrative example of the gTMT in action. When parental genotypes are missing, we propose to further extend gTMT by using Bayesian calculations on child genotypes to model parental genotypes as intermediate random variables.

## 1 Introduction

Genome-wide association studies (GWAS) are typically conducted on population-sampled individuals. Although GWAS identify statistically significant associations between genetic variants and phenotypes, such associations are not interpreted as causal findings due to possible confounding factors that are associated with both genotypes and phenotypes. This is not a weakness of GWAS, but rather it is a general limitation of observational studies where associations are identified.

In contrast, family-based studies sample closely related individuals based on pedigrees to estimate linkage between genetic variants and phenotypes of interest, while accounting for common confounding factors such as population structure and non-random mating [1–15]. However, formally establishing causal inference in family-based designs has received less attention. We have recently proposed a framework and robust test of causality in studies involving population-sampled parent-child trios, called the *Transmission Mean Test* (TMT) [16].

The TMT builds on the Neyman-Rubin *potential outcomes framework* — a gold standard statistical framework for causal inference [17–19] — based on the randomization of alleles transmitted during the meiosis process. The TMT is designed to have a general applicability to a broad class of phenotypes (continuous, count, or dichotomous traits). The TMT demonstrates robustness to a wide-range of confounding factors including population structure, parent effects, and non-genetic variation.

The original TMT is designed for trio studies that sample two parents and one child per family. However, many nuclear family datasets include multiple offspring per family. For example, the Norwegian Mother and Child Cohort Study (MoBa) includes approximately 16,400 families with two or more offspring in addition to around 44,000 parent-child trios [15, 20]. Families with multiple offspring are commonly observed among first-degree relatives in biobanks that originally intend to sample unrelated individuals. For example, previous work identified around 1,000 trios and 37 quartets (two parents and two offspring) in the UK Biobank cohort [21]. Additionally, the Finnish biobank FinnGen contains more than 12,000 nuclear families for which both parents and one or more offspring were directly genotyped [22, 23].

Here, we extend the TMT and its underlying causal inference framework to population-sampled nuclear families, where each family contains two parents and any number of offspring. The primary objective is to infer causal effects from genotype to phenotype when family genotypes are available and the phenotype has been measured on the offspring. We consider a sample of *J* nuclear families, each having two parents. Within family *j* (*j* ∈ [1 : *J*]), there are *K*_*j*_ offspring, *K*_*j*_ ∈ [1, 2, · · · ]. A special case of this is the trio study that samples two parents and one child per family so that *K*_*j*_ = 1 for all *j* ∈ [1 : *J*].

We consider the setting where unphased biallelic single nucleotide polymorphism (SNP) data have been collected on *I* SNPs per individual. For a SNP, let *a* and *b* be the two alleles that generate three possible genotypes {*aa, ab, bb*}. We numerically encode these three genotypes by {0, 1, 2}, respectively. In family *j*, let 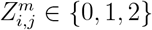 be genotype *i* of the mother, 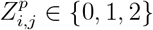 genotype *i* of the father, and *G*_*i,j,k*_ ∈ {0, 1, 2} genotype *i* of offspring *k* in family *j, k* ∈ [1 : *K*_*j*_], *j* ∈ [1 : *J*]. Figure S1 depicts a full set of genotypes in a nuclear family, as well as the inheritance process of randomized allele transmissions. We additionally denote 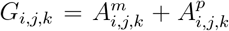 for each child, where 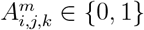 is the allele transmitted to offspring *k* in family *j* from the maternal side and 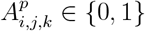 the allele from the paternal side (where 0 is *a* transmission and 1 is *b* transmission). Let *Y*_*j,k*_ be the phenotype of interest of offspring *k* in family *j*, where this phenotype can be real values, whole numbers, or dichotomous indicators {0, 1}. Writing ***Y*** = {*Y*_*j,k*_} and ***G***_*i*_ = {*G*_*i,j,k*_}, the goal is to infer the causal effect from ***G***_*i*_ to ***Y*** for *i* ∈ [1 : *I*]. We quantify this causal effect as the expected difference between potential outcomes of the traits corresponding to different alleles transmitted from parents to their offspring, i.e., what is typically called the *average causal effect* (ACE) from ***G***_*i*_ to ***Y*** . We develop an unbiased estimator of the ACE, together with its sampling variance estimate, to derive a test-statistic for testing the hypothesis of zero ACE versus non-zero ACE.

This paper introduces theoretical and technical innovations, building on previous work. First, to construct trait-level potential outcomes, we detail a model of the phenotype that accounts for within-family effects shared by siblings, forming the basis to quantify within-family covariances (among siblings) and between-family covariances (among offspring from different families). Second, we show how our proposed unbiased ACE estimate incorporates the phenotype values so that the estimate is robust to model misspecification and unmodeled polygenic background effects. Third, when estimating the sampling variance of the unbiased ACE estimator, we take into account covariances among all offspring to derive an estimate for the variance that shows the desired operating characteristics leading to a valid test-statistic. Along with our earlier work [16], we develop theory and methods for an atypical scenario in causal inference where two parallel randomized experiments of the same “treatment” — one per parent — are considered simultaneously on a set of subjects.

Our work has both connections to and differences with other methods in family-based studies. We previously showed the transmission disequilibrium test (TDT) [1–4] is a test of causality and a special case of our framework in ref. [16]. Nuclear family data have been utilized for association analyses [5–7, 12–14], but these existing approaches do not test for causality. Another line of research utilizes phased genomes of parent-child trios and requires assumptions about recombination rates to test for probabilistic independence based causality [24, 25], which is a different approach than that taken here.

The remainder of this paper starts by revisiting the TMT method for parent-child trios in Section 2.1. Then we establish the generalized TMT (gTMT) framework for nuclear families by developing an unbiased ACE estimator in Section 2.2, deriving its sampling variance estimate in Section 2.3, and proposing a hypothesis test in Section 2.4. We simulate nuclear family data to validate the performance of the gTMT in Section 3.1, contrast the gTMT with family-based association methods in Section 3.2, analyze UK Biobank data as an illustrative example in Section 3.3, and propose an approach to handle missing parental genotypes in Section 3.4. In Section 4, we develop and detail the underlying models and theory of the proposed framework to demonstrate that the gTMT is a valid causal inference method.

## 2 Proposed method

### 2.1 The TMT statistic for parent-child trios

We first review the previously proposed TMT for parent-child trios. The TMT is conducted on a per locus basis, so we drop the genetic locus index *i*. For parent-child trios, one child is observed per family so we drop the sibling subscript *k*. Let *N* be the total number of randomized allele transmissions; in trio studies, *N* equals the total number of heterozygous parents in that

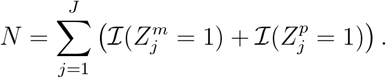

In ref. [16], we defined the TMT parameter *δ*_TMT_ as

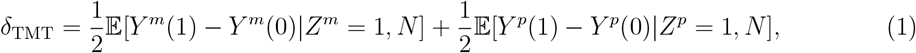

where *Y* (1) and *Y* (0) are the potential outcomes of the two possible transmitted alleles. We fully detail these quantities in Section 4 for the generalized setting considered here. We showed that *δ*_TMT_ is non-zero if and only if the ACE from the child genotype *G* to the child phenotype *Y* is non-zero (i.e., there exists a non-zero causal effect from *G* to *Y*). We constructed a hypothesis test where the null hypothesis is *H*_0_ : *δ*_TMT_ = 0 and the alternative is *H*_1_ : *δ*_TMT_ ≠ 0. We defined the TMT statistic for the hypothesis test as follows.

#### Definition 1

(TMT statistic for parent-child trios). *Define the assignment indicators*

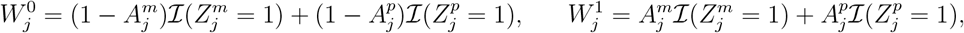

*and define the population mean estimate*

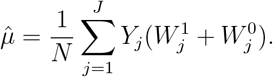

*The original TMT statistic is*

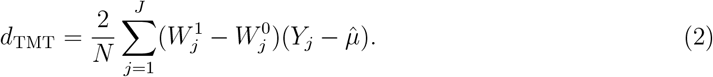

The assignment indicators capture whether *a* or *b* alleles were transmitted for each heterozygous parent of a child. In ref. [16], we proved that *d*_TMT_ is an unbiased estimator of *δ*_TMT_ in that 𝔼[*d*_TMT_|*N* ] = *δ*_TMT_. We developed a sampling variance estimate 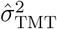 for 𝕍(*d*_TMT_|*N*) when *H*_0_ is true and is a lower bound for 𝕍(*d*_TMT_|*N*) when *H*_1_ is true. We defined the test statistic 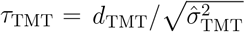 with the *p*-value calculated either by ℙ (|*X*| ≥ |*τ*_TMT_|) where *X* ∼ Normal(0, 1) or by permutation.

### 2.2 The gTMT statistic for nuclear families

Here, we allow each family to have an arbitrary number of offspring. Let *K*_*j*_ be the total number of offspring in family *j, K*_*j*_ ∈ [1, 2, 3, · · · ], *j* ∈ [1 : *J*]. Note that *K*_*j*_ = 1 in ref. [16] for parent-child trios. The total number of randomized allele transmissions *N* is now calculated as

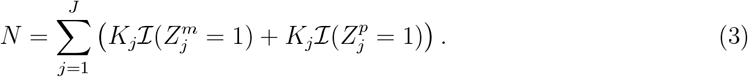

The gTMT parameter is the same as the original TMT parameter *δ*_TMT_ in Eq. (1). In Theorem 1, we show that *δ*_TMT_ is zero if and only if the ACE from the child genotype *G* to the child phenotype *Y* is zero, so that a non-zero *δ*_TMT_ indicates the existence of a causal effect. A hypothesis test can be conducted by testing the null hypothesis *H*_0_ : *δ*_TMT_ = 0 versus the alternative *H*_1_ : *δ*_TMT_ ≠ 0. To do so, we define the gTMT statistic *d*_gTMT_ as follows.

#### Definition 2

(gTMT statistic). *Define the assignment indicators*

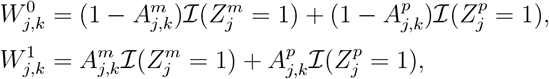

*and the population mean estimate*

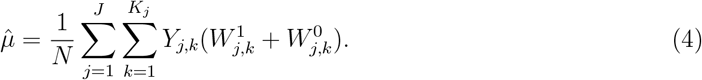

*The gTMT statistic is*

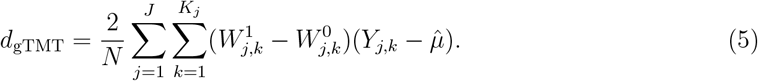

The factor 2*/N* comes from the fact that there is a 1*/*2 probability of ending of in the “control” or “treatment” groups for each transmission, so we divide the sum in Eq. (5) by *N/*2. In Theorem 2, we show that *d*_gTMT_ is an unbiased estimator for *δ*_TMT_ in that 𝔼[*d*_gTMT_|*N* ] = *δ*_TMT_. Figure 1 is a schematic showing how the randomized transmission of alleles leads to randomized assignments to the “control” and “treatment” groups, and the statistic *d*_gTMT_ measuring their difference.

**Figure 1:**
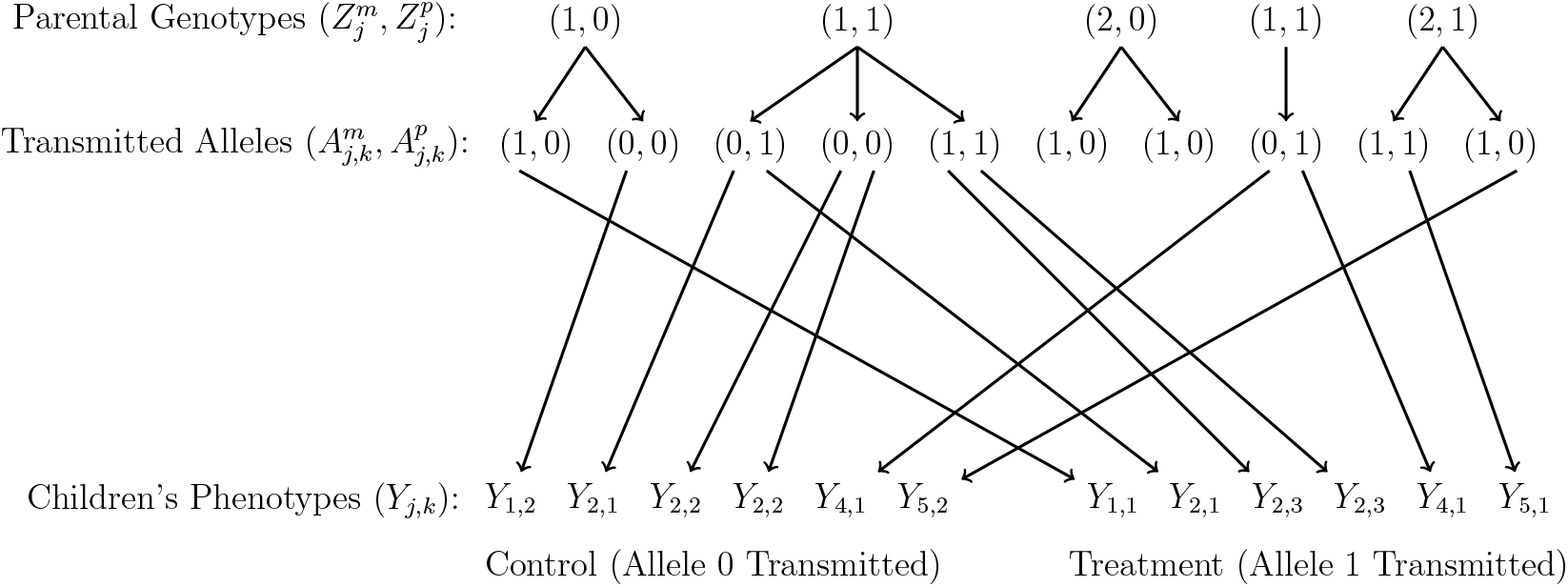
Schematic of the gTMT randomization framework. For offspring *k* in family *j*, if a heterozygous parent transmits an allele 0 to the child, the corresponding phenotype *Y*_*j,k*_ is included in the “control” group. If a heterozygous parent transmits an allele 1, *Y*_*j,k*_ is included in the “treatment” group. The top layer to the middle shows the allele transmission from parents to offspring. In the middle layer, the number of offspring per family varies. The bottom layer shows how the trait values are assigned to the control and treatment groups, the difference of which is measured by *d*_gTMT_.

### 2.3 Sampling variance of the gTMT statistic

As Figure 1 presents, one child can be assigned twice, once, or zero times to either the treatment group or the control group. The same child can also be assigned simultaneously once to each group, which will have zero contribution to *d*_gTMT_ based on Eq. (5) and does not contribute to the variance of *d*_gTMT_. These properties show how our particular setting is an atypical instance of causality under randomization (see ref. [16] for a full discussion). This leads to four groups contributing to the overall sampling variance of *d*_gTMT_, each of which may have a different group-specific variance:

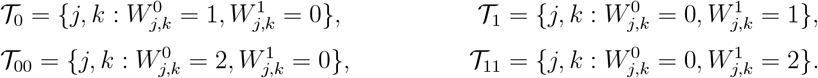

In 𝒯_0_, one heterozygous parent transmitted an allele 0 to the child, assigning the child phenotype to the “control”. In 𝒯_1_, one heterozygous parent transmitted an allele 1 to the child, assigning the child phenotype to the “treatment”. In 𝒯_00_, two heterozygous parents transmitted two alleles 0 to the child, assigning the child phenotype twice to the control. In 𝒯_11_, two heterozygous parents transmitted two alleles 1 to the child, assigning the child phenotype twice to the treatment. For each of these four sets, we form the following estimates:

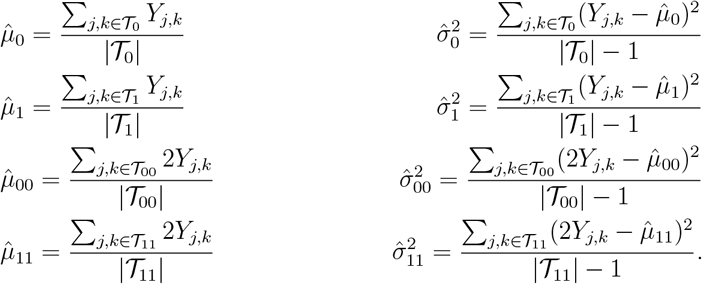

We form the sampling variance estimate for *d*_gTMT_ as:

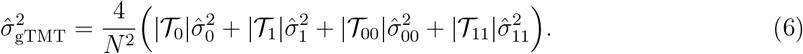

We discuss the theoretical foundations of the sampling variance estimate in Section 4.5, where we disentangle the variances and covariances within and between families.

### 2.4 Proposed hypothesis test

We form the following test statistic

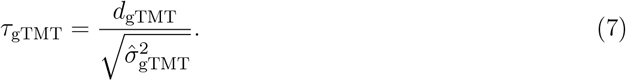

By the Central Limit Theorem, *τ*_gTMT_ is approximately Normal(0, 1) when the null hypothesis of no causality is true (see *Remark* in Appendix A.6). In this case, the *p*-value is calculated by *p*_gTMT_ = ℙ (|*X*| ≥ |*τ*_gTMT_|) where *X* ∼ Normal(0, 1). One can also use a permutation null distribution as an alternative to the Normal(0, 1) distribution to calculate *p*-values (details in Appendix A.1). We verify that the permutation null and the Normal(0,1) yield similar results in Section 3.1.

## 3 Results

### 3.1 Performance of gTMT as a test of causality

We implemented an algorithm (detailed in Appendix B.1) to simulate genotypes of 3,000 nuclear families including 1,000 trios (two parents and one child), 1,000 tetrads (two parents and two offspring) and 1,000 quintets (two parents and three offspring). This produced 6,000 offspring and 6,000 parents with 100,000 SNPs per individual. As described in Appendix B.1, we simulated tractable population structure by generating genotypes via a standard admixture model [26– 29]. We randomly assigned 100 SNPs to be causal and followed Appendix B.2 to generate child phenotypes across various levels of true heritability *h*^2^.

#### Unbiased estimation of the target parameter

We confirmed the accuracy of *d*_gTMT_ as an unbiased estimate of the TMT parameter *δ*_TMT_ among various levels of *h*^2^ (Figure 2A).

**Figure 2:**
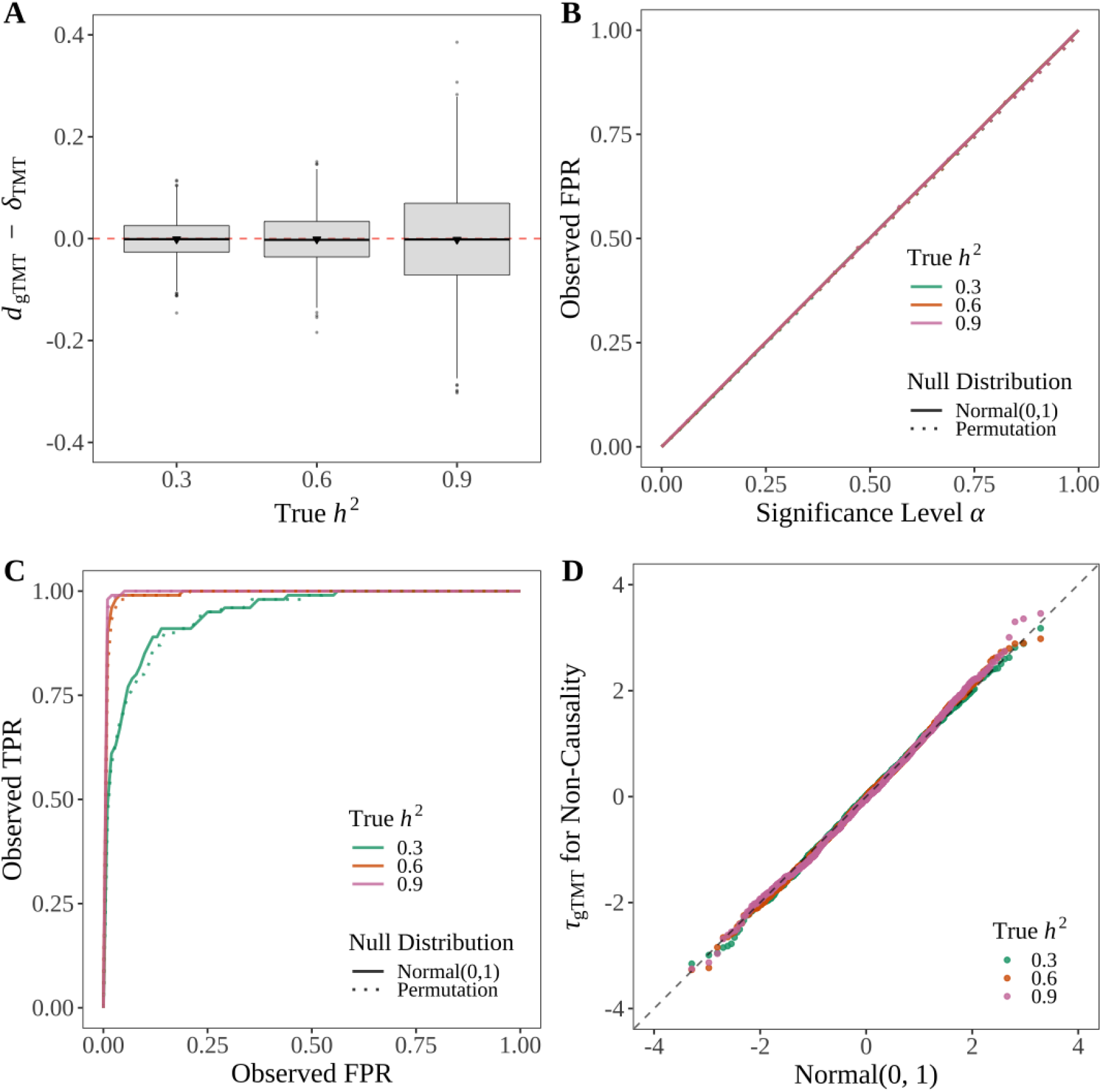
gTMT is a valid test of causality for population-sampled nuclear families. **(A)** *d*_gTMT_ is an unbiased estimator of *δ*_TMT_. We generated 1,000 family trios, 1,000 tetrads and 1,000 quintets (in total 3,000 families with 6,000 offspring) from a structured population (*F*_ST_ = 0.2) with 100,000 SNPs per individual and we randomly chose 100 genetic loci to be causal. We simulated the offspring phenotypes for *h*^2^ ∈ { 0.3, 0.6, 0.9 } . Shown are 1,000 randomly chosen instances of *d*_gTMT_ − *δ*_TMT_ for each *h*^2^. **(B)** gTMT controls FPR at the desired significance level per setting. **(C)** The ROC curve from a randomly chosen simulated data set. **(D)** The distribution of the test statistic *τ*_gTMT_ is well approximated by Normal(0,1) when the null hypothesis of no causality is true.

#### Causal FPR and statistical power

We tested for non-zero ACE per locus and calculated *p*-values for both causal and non-causal SNPs by utilizing both the Normal(0, 1) null distribution and the permutation null. We then computed the false positive rate (FPR) and the true positive rate (TPR) at various significance thresholds in the unit interval [0, 1]. The gTMT test controlled the FPR at the desired significance level across all levels of *h*^2^ (Figure 2B). The receiver operating characteristic (ROC) curve demonstrated gTMT having statistical power for detecting direct causal effects for all levels of *h*^2^ (Figure 2C).

#### Accuracy of the null distribution

We confirmed that the distribution of the test statistic *τ*_gTMT_ is well approximated by the standard Normal (0, 1) distribution when the null hypothesis of no causality is true (Figure 2D).

### 3.2 The gTMT versus family-based association methods in the presence of confounding

We compared gTMT with association methods applicable to family data, noting that those methods are intended for associations. The goal of the comparison is to show that gTMT is accurate for identifying causal effects in contrast to these association methods when the causal effects are confounded.

A popular association method is the family-based association test (FBAT) [5–8], which is a score test based on the FBAT statistic 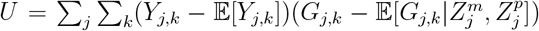. Conditioning on the trait *Y* and parental genotypes, FBAT derives the variance of *U* by 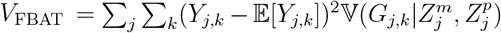 and calculates the test statistic 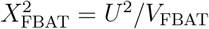, with 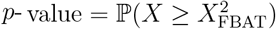 where *X* has a 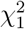 distribution. Another approach called the family-based GWAS (FGWAS) builds on a regression model 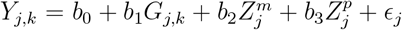 to estimate *b*_1_ as the association parameter between *Y* and *G* while controlling for parental genotypes *Z*^*m*^ and *Z*^*p*^ [12–14]. Distinctions between FGWAS and the TMT have been further detailed in ref. [16]; it has also been shown that a variety of confounding effects can impact the performance of FGWAS [30].

Here, we considered family-specific confounding effects that are correlated with the parental genotypes. We evaluated the statistical power of gTMT, FBAT and FGWAS in detecting causal genetic variants. Note that associations can be affected by confounding, whereas a causal inference method should be robust to such confounding. We implemented an algorithm (detailed in Appendix B.1) to simulate genotypes of 1,500 nuclear families (500 trios, 500 tetrads and 500 quintets). This produced 3,000 offspring and 3,000 parents with 100,000 SNPs per individual. We randomly chose 100 causal SNPs and generated child traits by

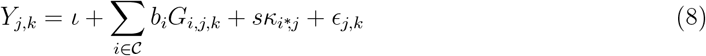

where 𝒞 is the set of causal SNPs and 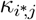 is the family effect correlated with parental genotypes at a causal locus *i*^∗^ randomly chosen from 𝒞 (details in Appendix B.3). We applied gTMT, FBAT and FGWAS to compute *p*-values at the causal loci. Specifically, for 3,000 iterations, we generated child genotypes at the causal loci, simulated child traits by Eq. (8), and applied the three methods to derive *p*-values. This produced 3,000 p-values per causal locus per method. We calculated empirically the type II error (*β*) as the proportion of *p*-value greater or equal to a significance-level *α* = 0.05 among 3,000 *p*-values for each method, i.e., *β* = (no. *p*-values ≥ *α*)*/*3000. Then we used 1 − *β* to estimate the statistical power.

Figure 3 shows the effect of confounding from the causal locus *i*^∗^. We considered a range of confounding effect sizes *s* (in Eq. (8)) from *s* = 0 (no confounding) to *s* = 5 and computed the statistical power for each method. The gTMT maintained the same power across all levels of the confounding effect size, while both FBAT and FGWAS had decreasing power when the confounding effect size increased. We conducted the same power analysis for larger samples: (i) 6,000 offspring from 3,000 families (1,000 trios, 1,000 tetrads and 1,000 quintets); and (ii) 12,000 offspring from 6,000 families (2,000 trios, 2,000 tetrads and 2,000 quintets). Both scenarios showed higher power for the gTMT than the other two methods in the presence of confounding effects (Figure 3).

**Figure 3:**
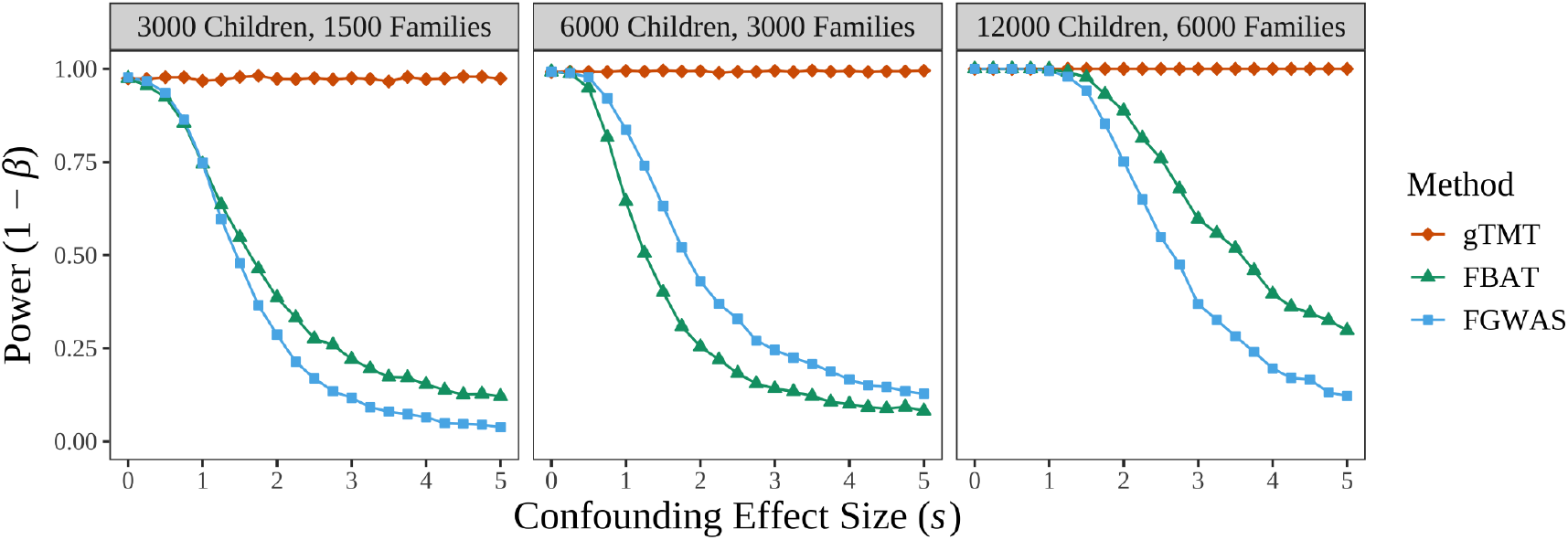
Statistical power for detecting causal effects of the generalized TMT (gTMT), family-based association test (FBAT), and family-based GWAS (FGWAS) in the presence of confounding.

### 3.3 UK Biobank data analysis

#### Identifying nuclear families

We used kinship estimates from `KING` [31], age, and gender to identify 990 trios (two parents and one child) and 37 quartets (two parents and two offspring) with 1,064 distinct offspring and 2,054 parents in total. We detail the procedure for identifying nuclear families in Appendix C.1.

#### The gTMT for blood pressure

We analyzed three blood pressure measures, including systolic blood pressure (SBP), diastolic blood pressure (DBP), and pulse pressure (PP) where PP = SBP − DBP. We detail the quality control pipeline for these blood pressure measures in Appendix C.2.

We implemented gTMT across 9,563,967 autosomal SNPs that have minor allele frequency (MAF) greater than 1% and imputation information scores [32] greater than 0.8. We calculated one *p*-value per SNP and display − log_10_(*p*) in Figure 4 as the genome-wide gTMT profile. For all three blood pressure measures, we successfully detected several genome-wide significant signals with *p*-value *<* 5 *×* 10^−8^.

**Figure 4:**
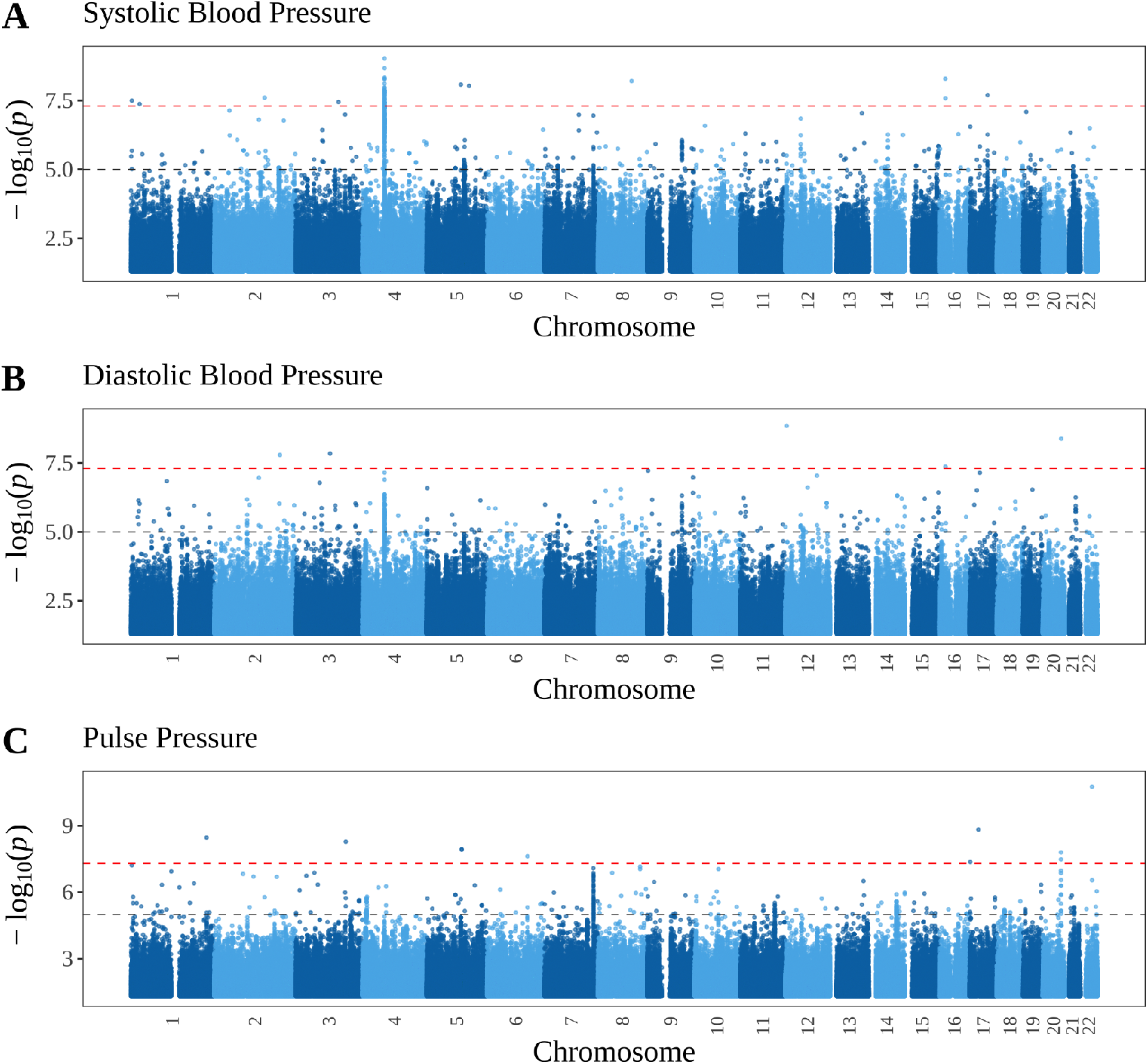
UK Biobank genome-wide gTMT profile for blood pressure measures including (A) systolic blood pressure; (B) diastolic blood pressure; (C) pulse pressure. The gray dashed line is for *p*-value= 1 *×* 10^−5^. The red dashed line is for *p*-value= 5 *×* 10^−8^.

#### Comparing to family-based association methods

We conducted FBAT and FGWAS to detect associations between autosomal SNPs and blood pressure measures. Figure 5 displays quantile-quantile plots comparing *p*-values from gTMT, FBAT, and FGWAS versus *p*-values under the null hypothesis of no causality that are Uniform(0, 1) distributed. In general, both FBAT and FGWAS were more conservative than gTMT. For the pulse pressure (PP) measure, only gTMT successfully detected genome-wide significant signals while both FBAT and FGWAS did not. One possible reason is that intra-individual random effects acting as confounders are driving the FBAT and FGWAS statistical significance in SBP and DBP. When analyzing PP, since PP = SBP − DBP, intra-individual random effects are likely eliminated. For PP, therefore, statistical significance was absent for FBAT and FGWAS, while gTMT maintained notable statistical significance.

**Figure 5:**
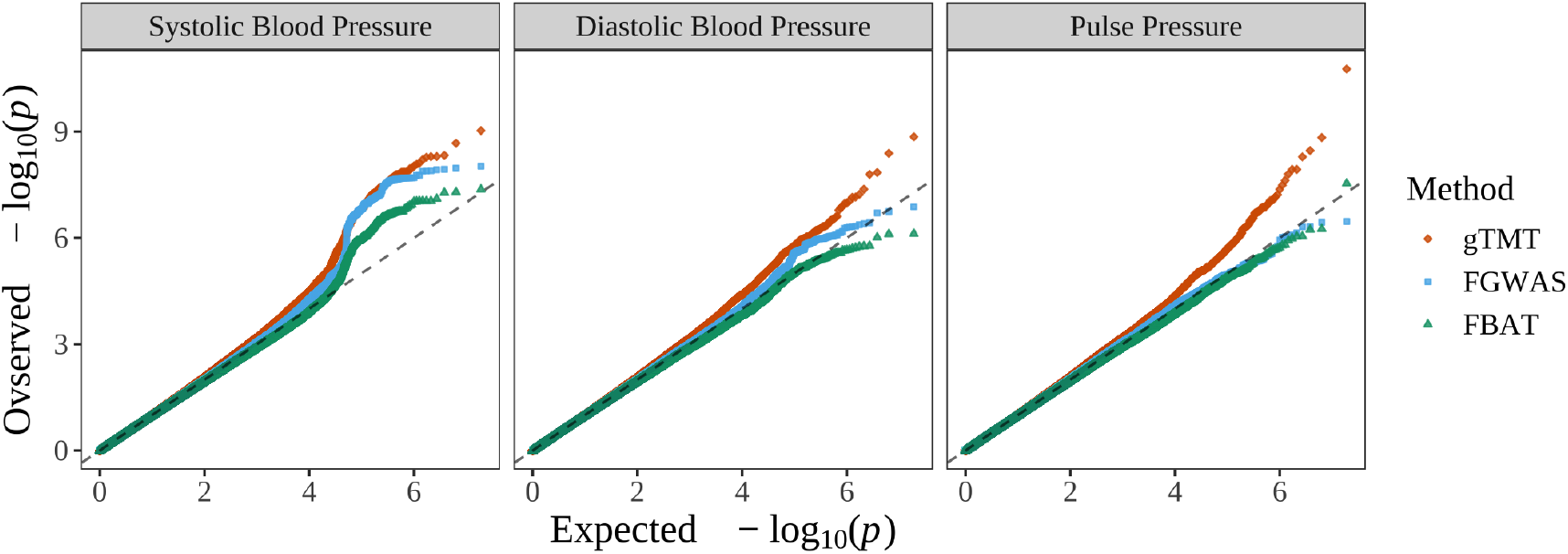
Quantile-quantile plots of observed *p*-values from gTMT, FBAT, and FGWAS versus expected *p*-values under the null hypothesis of no causality. Note that *p*-values above the identity line show signal for statistical significance.

### 3.4 Missing parental gentoypes

When one or more parental genotype is missing, the following is a possible strategy to extend the gTMT by using Bayesian probabilities to model parental genotypes as intermediate random variables. Let {*G*_*j,k*_} be the genotypes of all offspring in family *j*. By using Bayes theorem, the conditional probability for observing the possible parental genotype pairs is

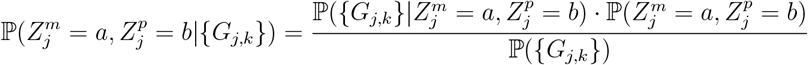

where *a, b* ∈ {0, 1, 2}. For the conditional probability 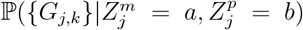, we follow Mendel’s laws to calculate 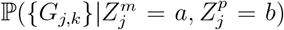 among family trios (Table S1) and tetrads (Table S2). In an analogous way, one can enumerate 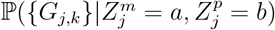 for families with an arbitrary number of offspring. The other two probabilities 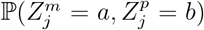 and ℙ({*G*_*j,k*_}) can be calculated empirically based on the data in the study or other data sets sampled from the same population. This pipeline could be directly applied to sibling studies where parental genotypes are generally unavailable.

## 4 Theoretical foundations of the gTMT

We develop models and theory here to demonstrate that the gTMT is a valid test of causality. The following is an extension of the potential outcomes framework for trios in ref. [16] to the scenario we consider here where families may have more than one offspring.

### 4.1 The trait model

The trait for offspring *k* in family *j* is modeled as:

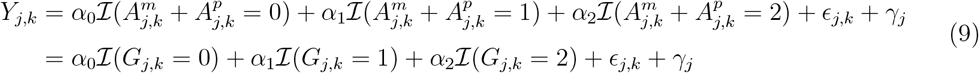

where *γ*_*j*_ is a random variable shared by siblings in family *j*, while *ϵ*_*j,k*_ is a random variable specific to offspring *k* in family *j*. We do not make specific assumptions about the expected values of *ϵ*_*j,k*_ and *γ*_*j*_ nor do we require exogeneity. The dependence structure of these random variables is articulated in Assumption 2.

Here we retain Assumption 1 from ref. [16] that the genetic effects are either non-decreasing or non-increasing.

#### Assumption 1.

*The conditional expectation of offspring phenotype given parent-transmitted alleles*, 𝔼 [*Y* |*A*^*m*^, *A*^*p*^], *is either a non-decreasing function of A*^*m*^ + *A*^*p*^ *such that*

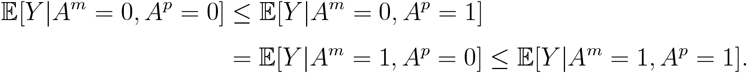

*or* 𝔼 [*Y* |*A*^*m*^, *A*^*p*^] *is analogously a non-increasing function of A*^*m*^ + *A*^*p*^.

Note that under the trait model in Eq. (9), if 𝔼 [*Y* |*A*^*m*^, *A*^*p*^] is non-decreasing, then *α*_0_ ≤ *α*_1_ ≤ *α*_2_; if 𝔼 [*Y* |*A*^*m*^, *A*^*p*^] is non-increasing, then *α*_0_ ≥ *α*_1_ ≥ *α*_2_.

### 4.2 Potential outcomes and causal effects

In order to characterize the causal effect of the parent-transmitted alleles (*A*^*m*^, *A*^*p*^) on the offspring phenotype *Y*, we previously [16] modeled potential outcomes for *Y* in terms of each parent individually. We formulated variables representing the phenotype that the offspring would have developed if receiving a particular allele from the parent. Let *A* be the parent-transmitted allele. The potential outcomes are *Y* (*A* = 0) and *Y* (*A* = 1), which we sometimes simplify as *Y* (1) and *Y* (0) when there is no ambiguity. Then the observed trait value *Y* can be written as a function of the potential outcomes,

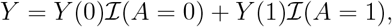

where ℐ (·) is the indicator function. When considering a particular parent’s contribution, let *Y* ^*m*^(0) and *Y* ^*m*^(1) be potential outcomes for the maternal side and *Y* ^*p*^(0) and *Y* ^*p*^(1) for the paternal side. Here, we extend this to potential outcomes in the scenario of nuclear families with an arbitrary number of offspring per family. Under the trait model in Eq. (9), for offspring *k* in family *j*, the potential outcomes are

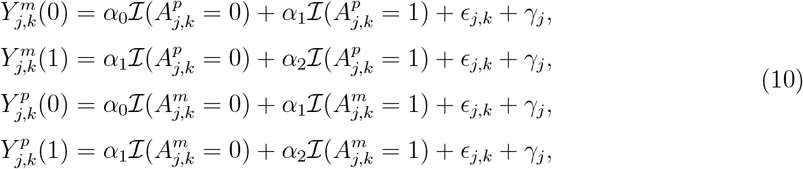

leading to two pairs of causal effects per parent:

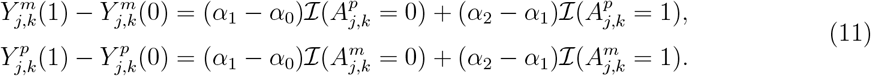

We use expectations of these causal effects to define the average causal effect (ACE) as follows; this definition applies to all offspring, so we omit the family and sibling indices *j, k* for the rest of this subsection.

#### Definition 3

(ACE for an allele). *The ACE of a parent-transmitted allele A on offspring trait Y is the average difference between potential outcomes Y* (*A* = 1) *and Y* (*A* = 0): ACE(*A* → *Y*) = 𝔼 [*Y* (*A* = 1)] − 𝔼 [*Y* (*A* = 0)]. *We say A is directly causal for Y, denoted A* → *Y, if* ACE(*A* → *Y*)≠ 0.

We derive ACE per parent explicitly as:

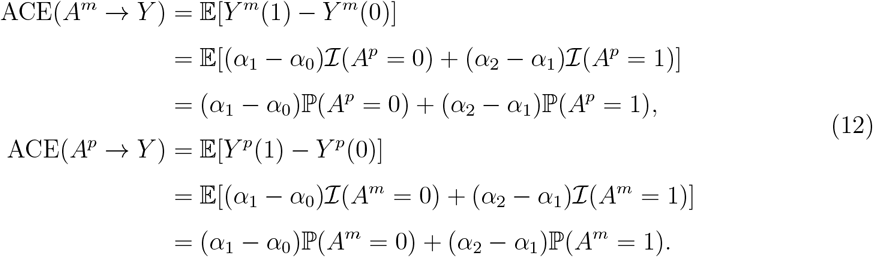

Assuming 0 *<* ℙ (*A*^*m*^ = 1) *<* 1 and 0 *<* ℙ (*A*^*p*^ = 1) *<* 1 in the sampled population, then ACE(*A*^*m*^ → *Y*) = ACE(*A*^*p*^ → *Y*) = 0 if and only if *α*_0_ = *α*_1_ = *α*_2_. This leads to the following definition of the ACE of the child genotype *G* = *A*^*m*^ + *A*^*p*^ on the child phenotype *Y*, denoted as ACE(*G* → *Y*).

#### Definition 4

(ACE for genotype). *Under the trait model in Eq*. (9), ACE(*G* → *Y*) = 0 *if and only if α*_0_ = *α*_1_ = *α*_2_. *Otherwise*, ACE(*G* → *Y*)≠ 0. *The offspring genotype G is directly causal for Y, denoted by G* → *Y, if* ACE(*G* → *Y*)≠ 0.

Based on Eq. (12) and Definition 4, it is trivial that the following three properties are equivalent:

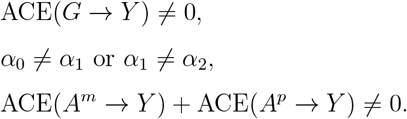

### 4.3 The gTMT parameter quantifies the causal effect

Recall that *N* is the total number of randomized allele transmissions from a parent to a child among all families, as previously defined in Eq. (3). The TMT parameter in Eq. (1),

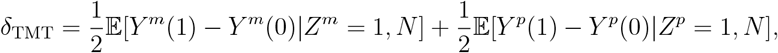

is the same value regardless of whether we are considering trios or generalized nuclear families since it is defined in terms of one parent and one child. We show that a non-zero value of δ_TMT_ implies non-zero ACE through the following theorem.

#### Theorem 1.

*Under the trait model in Eq*. (9) *and Assumption 1, the TMT parameter* δ_TMT_ = 0 *if and only if* ACE(*G* → *Y*) = 0.

We prove Theorem 1 in Appendix A.2.

### 4.4 The gTMT statistic is an unbiased estimate

In Section 2.2, we proposed *d*_gTMT_ as an unbiased estimate of δ_TMT_ in that 𝔼 [*d*_gTMT_|*N* ] = δ_TMT_, which we prove here. By Mendel’s Law of Segregation, the allele transmitted from a heterozygous parent to offspring is ℙ (*A*_*j,k*_ = *a*|*Z*_*j*_ = 1) = 1*/*2 for *a* ∈ {0, 1}. Since this randomization in meiosis precedes other factors such as the offspring individual effect *ϵ*_*j,k*_ and the family effect *γ*_*j*_ in Eq. (9), we make the following assumption.

#### Assumption 2.

*Under the trait model in* *Eq*. (9),

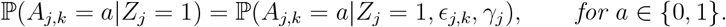

When Assumption 2 is satisfied, the following lemma holds, which will later be used to show that *d*_gTMT_ is an unbiased estimate of δ_TMT_.

#### Lemma 1.

*Under the trait model in Eq*. (9) *and given Assumption 2, the potential outcomes, Y*_*j,k*_(0) *and Y*_*j,k*_(1), *are conditionally independent of the parent-transmitted allele, A*_*j,k*_, *given a heterozygous parental genotype, Z*_*j*_ = 1, *written as*

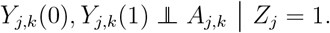

We prove Lemma 1 in Appendix A.3, and we later use Lemma 1 to prove Theorem 2.. Lemma 1 satisfies the unconfoundedness assumption in “intention-to-treat analysis” [19], which is the basic assumption in classical causal inference analysis under the potential outcomes framework.

Here we prove that 𝔼 [*d*_gTMT_|*N* ] = δ_TMT_ in Theorem 2. The proof is based on the trait model in Eq. (9) and requires Assumption 1 and Assumption 2. We show the following lemma as an intermediate step.

#### Lemma 2.

*For the generalized TMT statistic d*_gTMT_ *and the population-mean estimate* 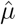,

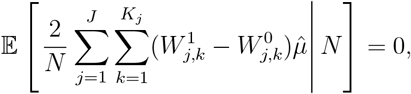

*so that*

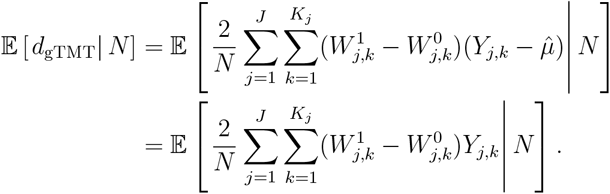

We prove Lemma 2 in Appendix A.4. Lemma 2 leads to the following theorem that states *d*_gTMT_ is an unbiased estimate of δ_TMT_.

#### Theorem 2.

*Under the trait model in Eq*. (9), *when Assumption 1 and Assumption 2 are satisfied, the generalized TMT statistic is unbiased for the TMT parameter in that*

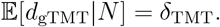

We prove Theorem 2 in Appendix A.5. To test the null hypothesis *H*_0_ : δ_TMT_ = 0 versus the alternative *H*_1_ : δ_TMT_≠ 0, we propose a sampling variance estimate for *d*_gTMT_ in Eq. (6) in the next section.

### 4.5 Estimate of the sampling variance

Here, we derive 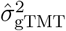 from Eq. (6) as an estimate of the sampling variance of *d*_gTMT_. Since the statistic *d*_gTMT_ contains a population mean estimate 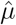, we first calculate the variance of the following statistic with known *µ*:

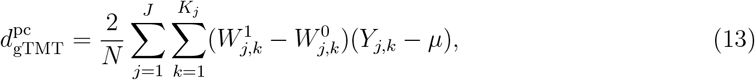

where

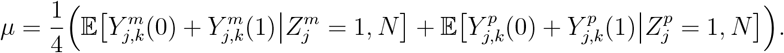

#### Theorem 3.

*When the null hypothesis that* δ_TMT_ = 0 *is true*,

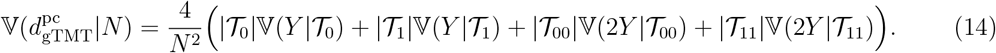

In general,

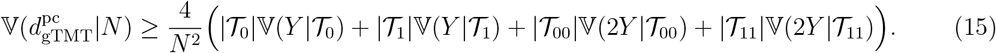

The proof of Theorem 3 is in Appendix A.6. This proof shows how to disentangle the variances and covariances within and between families. Defining 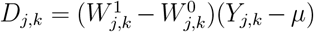 and referring back to the definition of 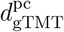 in Eq. (13), we can see that

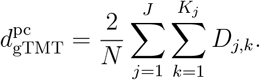

Notice that 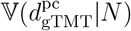 contains the covariance between siblings within the same family, i.e., ℂ (*D*_*j,k*_, *D*_*j,h*_|*N*) where *k* ≠ *h*. Also, 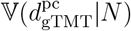 involves the covariance between offspring from different families, i.e., ℂ (*D*_*j,k*_, *D*_*l,s*_|*N*) where *j l*. In the proof of Theorem 3, we show that when the null hypothesis is true ℂ (*D*_*j,k*_, *D*_*j,h*_|*N*) = 0 and in general ℂ (*D*_*j,k*_, *D*_*j,h*_|*N*) ≥ 0. We also show that when the null hypothesis is true ℂ (*D*_*j,k*_, *D*_*l,s*_|*N*) = 0 and in general ℂ (*D*_*j,k*_, *D*_*l,s*_|*N*) ≥ 0. All these covariance properties are key to Theorem 3. Motivated by Eq. (14) in Theorem 3, we form 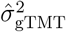 in Eq. (6) to estimate the variance of *d*_gTMT_ under the null hypothesis. Our simulations in Section 3.1 and Figure 2 empirically validate 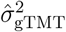 as an estimate of the variance of *d*_gTMT_ under the null hypothesis.

## 5 Discussion

Building on our earlier work for parent-child trios [16], we have extended this framework to population-sampled nuclear families, establishing a robust inference method to detect causal effects of genotype on phenotype. We have presented theory and methods to rigorously prove the intuition that family-based designs enable causal inference, owing to their inherent randomization through allele transmission and their observed robustness to common confounding effects such as population structure and non-random mating.

For studies where individuals are randomly sampled from a population, association tests and downstream methods such as fine-mapping have been established as a viable approach to identifying associations between genotype and phenotype. GWAS usually implement a linear mixed model regression to identify associations. Such regression approaches make the potentially strong assumption of exogeneity in that the non-genetic variation has zero covariance with the genetic variation. This assumption is not always satisfied for population-sampled nuclear families because family specific effects — *γ*_*j*_ in Eq. (9) — can introduce non-genetic factors that induce confounding between the child’s genotypes and phenotype. The existence of such confounding factors can generate false positives for GWAS results when including nuclear families. The gTMT is robust to these confounding factors because randomization has been leveraged, captured by the potential outcomes in Eq. (10) to derive the causal effects in Eq. (11).

For very large sample size population-based studies, there are often first degree relatives present. For example, we identified 28,895 candidate parent-child relationships in the UK Biobank data. One could use the gTMT and future extensions in conjunction with existing GWAS methods in these studies. The full set of related individuals could be set aside, while an association analysis is then applied to the remaining unrelated individuals. A set of SNPs showing associations in this first stage analysis could then be tested for causality using the gTMT in the related individuals that have been set aside. This strategy would increase the power of the secondary gTMT stage because fewer SNPs would be tested, and it would also provide a valid test of causality of the associated SNPs from the first stage.

Going beyond nuclear families to more general closely related individuals, future work could develop a framework where one can first probabilistically model missing parent genotypes and then conduct the gTMT. Large-scale family-based genetic data can have first degree relatives across multiple generations, such as offspring, parents, and grandparents. An example is the Framingham Heart Study that has recruited 5,209, 5,124, and 4,095 participants from three generations of residents in Framingham, with each generation as the offspring of the previous generation [33–35]. For such data, the gTMT could be implemented to analyze these data across generations.

The gTMT is currently designed to analyze common genetic variants. For rare genetic variants, the proportion of heterozygous parents would be small, affecting the total number of randomized allele transmissions from heterozygous parents and the power of the gTMT. This limitation has also been recognized for family-based association tests. Previous work has proposed methods to detect associations between rare variants and traits [11, 36–38]. An extension of the TDT for rare variant association tests has been developed for binary traits, but currently shows some inflated type I error rates [39]. An interesting future direction for the gTMT would be to build on these existing ideas to modify the gTMT to achieve reasonable power when analyzing rare variants.

## Resources

Reproducible documents are available at https://github.com/StoreyLab/causal-nuclear-family. An R package `geneticTMT` that implements the proposed method is available at https://github.com/StoreyLab/geneticTMT.

## Acknowledgments

This work was supported in part by United States National Institutes of Health grant R01 HG006448 to JDS. This research has been conducted using the UK Biobank Resource under Application Number 104628.

## Appendices

### A Theory

#### A.1 Permutation test

The permutation test is carried out by permuting the offspring phenotypes over *B* iterations. In each round of permutation, let 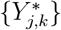 be the permuted phenotypes We calculate

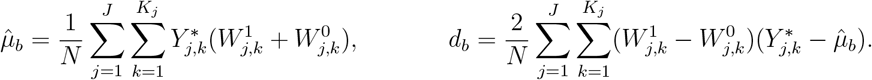

We then calculate the permutation p-value by 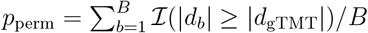.

#### A.2 Proof of Theorem 1

Plugging Eq. (11) into *δ*_TMT_ leads to the following.

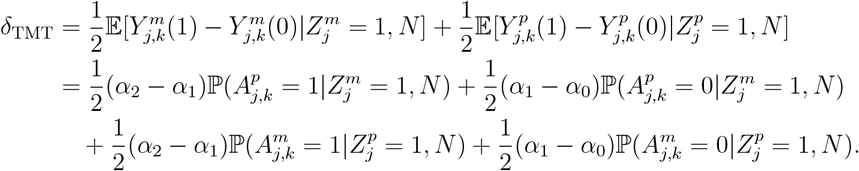

In practice 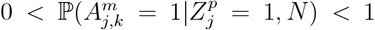 and 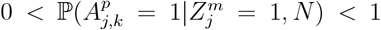. Under Assumption 1, either *α*_0_ ≤ *α*_1_ ≤ *α*_2_ or *α*_0_ ≥ *α*_1_ ≥ *α*_2_. Then *δ*_TMT_ = 0 if and only if *α*_0_ = *α*_1_ = *α*_2_, which implies ACE(*G* → *Y*) = 0 by Definition 4.

#### A.3 Proof of Lemma 1

We show on the maternal side 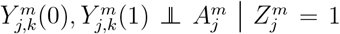. The proof for the paternal side follows the same strategy We first show that

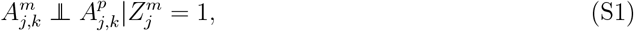

which is equivalent to prove

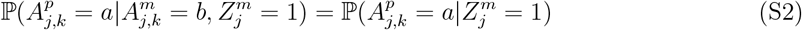

for *a, b* ∈ {0, 1}. First, we directly calculate the following conditional covariance:

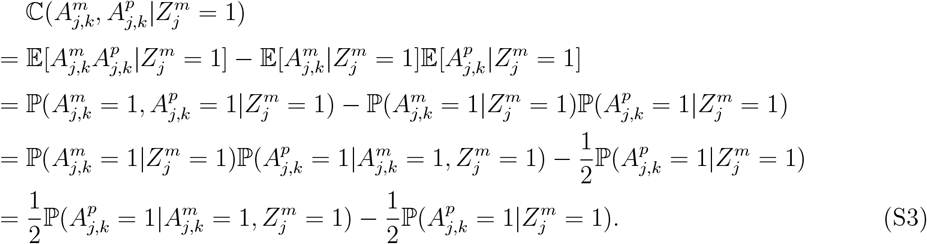

By the Law of Total Covariance,

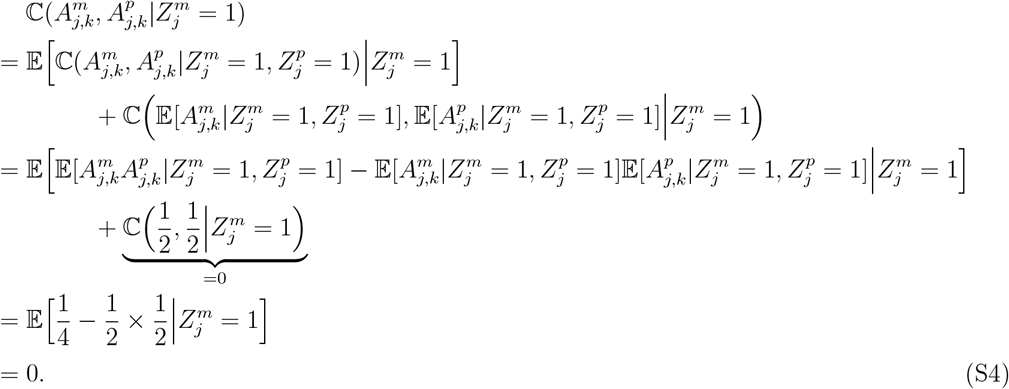

Since Eq. (S3) and Eq. (S4) are equal, this implies 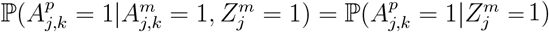 . Similarly, 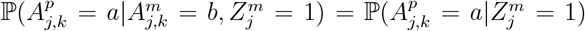 for *a, b* ∈ {0, 1}. Therefore, 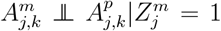. By Assumption 2, *ϵ*_*j,k*_ and *γ*_*j*_ are also independent of 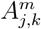 conditional on 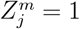. Thus, all random variables contributing to 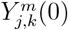 and 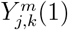 in Eq. (10) are independent of 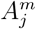 conditional on 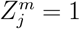, which completes the proof.

#### A.4 Proof of Lemma 2

We first show that 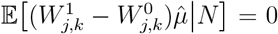 . We write 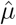 in Eq. (4) as 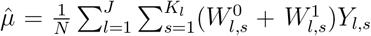 . Then

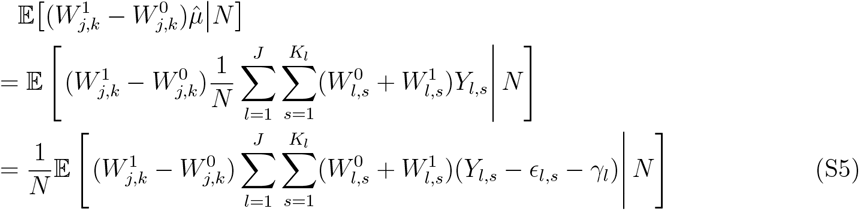

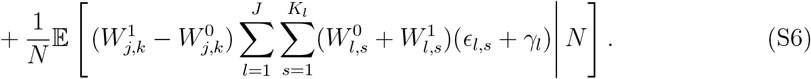

To show that Eq. (S5) equals 0, define

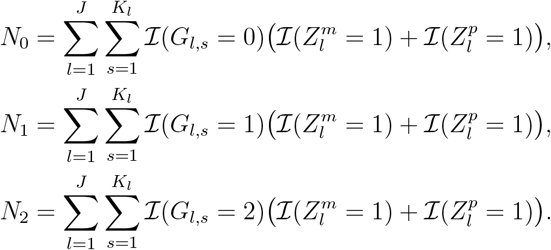

Also observe that

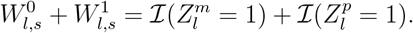

We use the above equations for *N*_0_, *N*_1_, *N*_2_, and 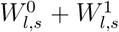. We also recall the trait model in Eq. (9) to derive Eq. (S5) as

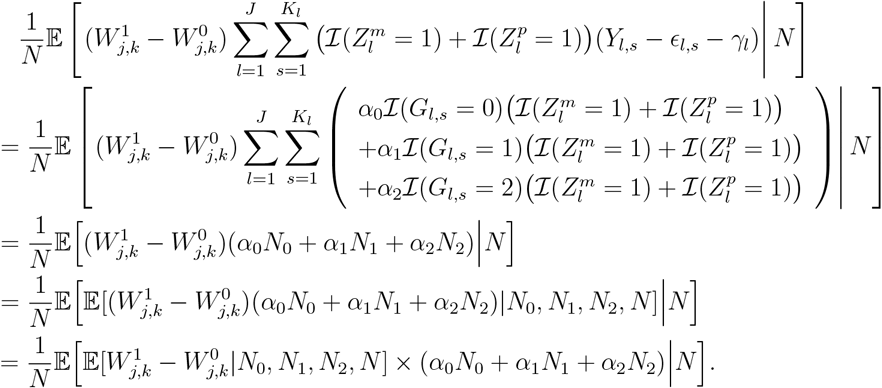

Note that

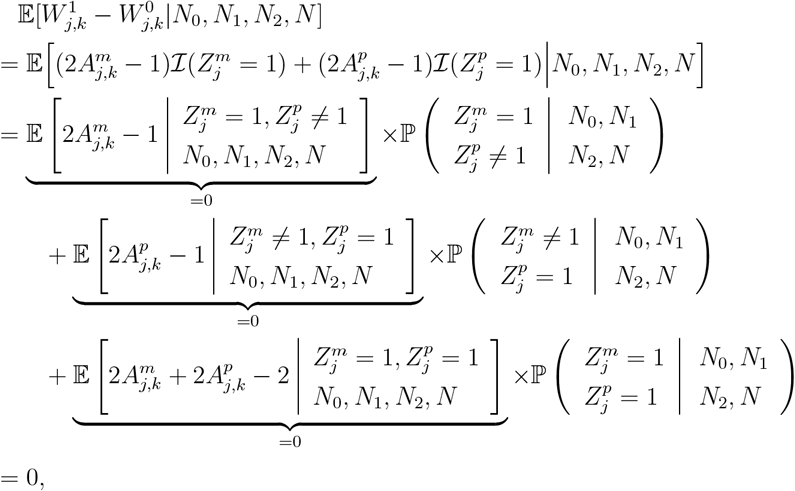

and therefore,

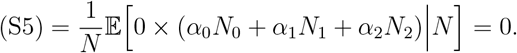

For Eq. (S6),

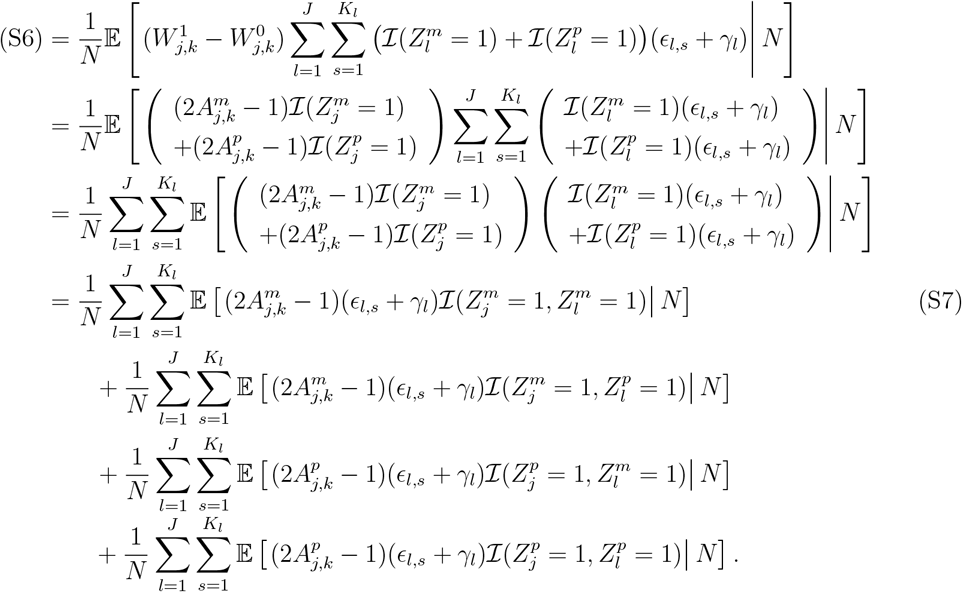

Now we show Line (S7) equals 0, and then the three lines that follow equal 0 by the same proof. By Mendel’s Law of Segregation,

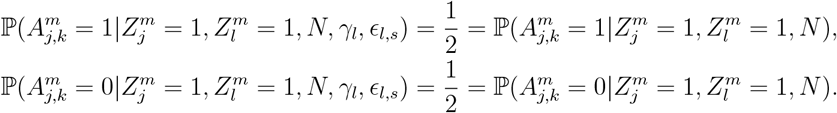

Then 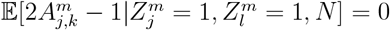 and 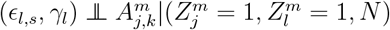 so that

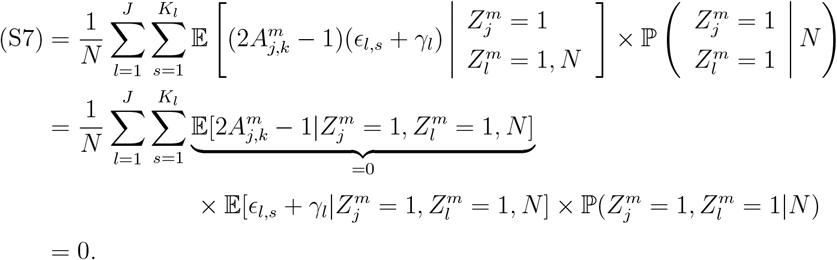

Eq. (S6) equals 0 and

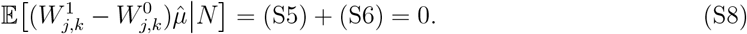

Therefore,

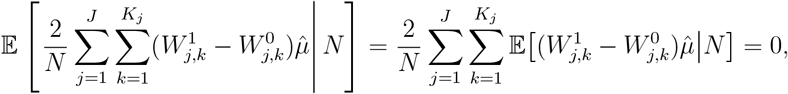

and

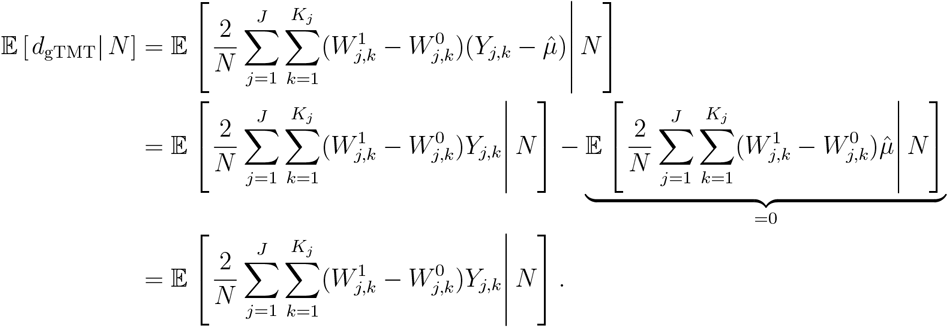

#### A.5 Proof of Theorem 2

Let 𝒥 be the total number of offspring in that 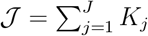.

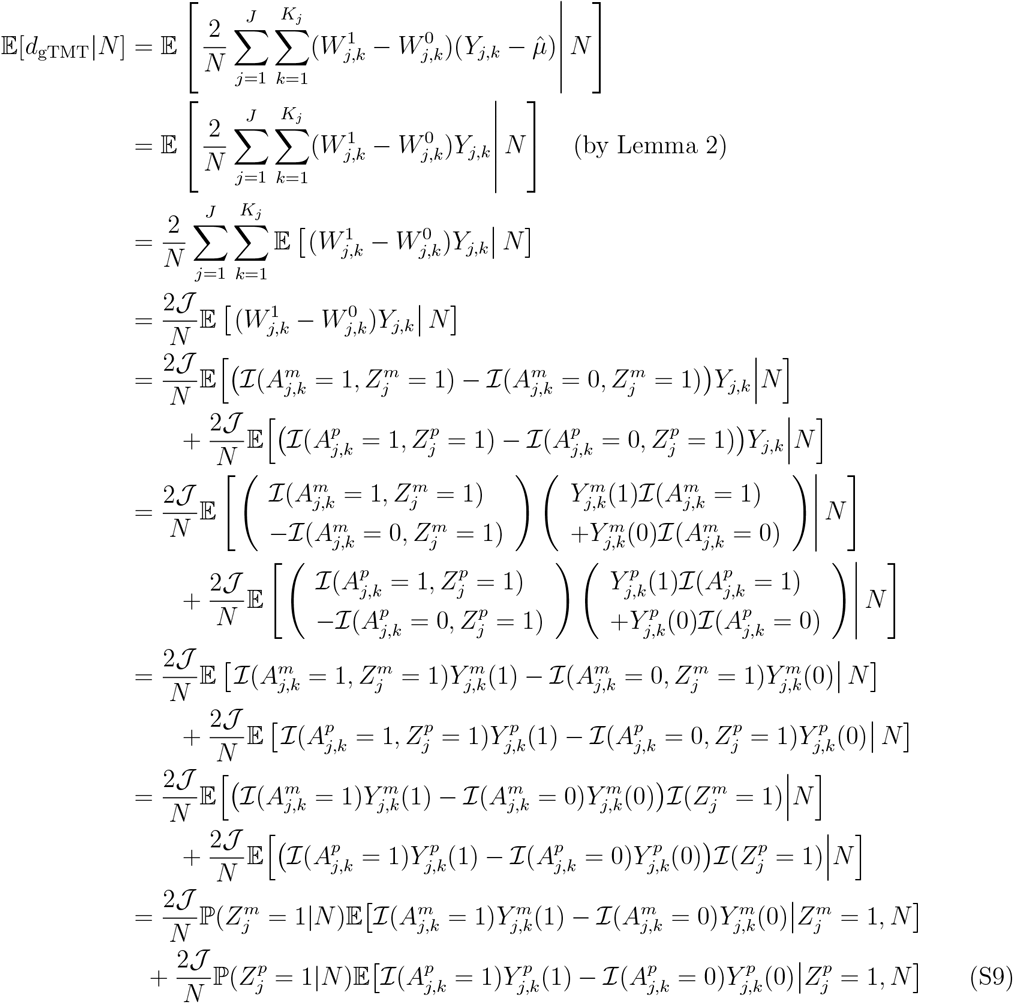

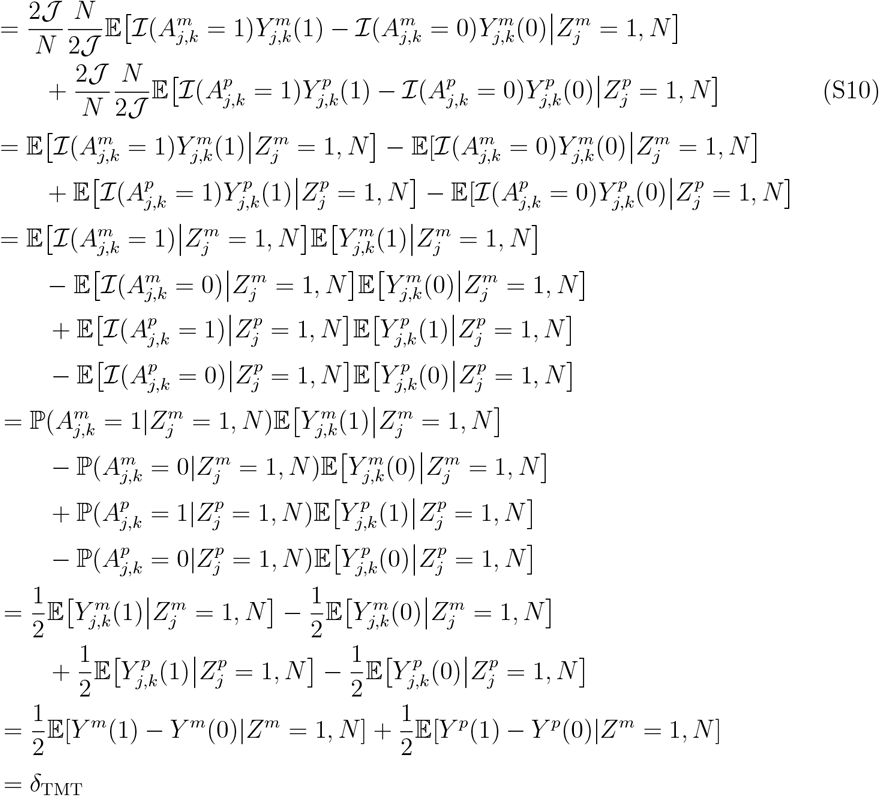

Eq. (S9) equals Eq. (S10) because ℙ(*Z*^*m*^ = 1|*N*) = ℙ(*Z*^*p*^ = 1|*N*) = *N/*(2𝒥), which follows because there are *N* parent-offspring pairs with heterozygous parental genotype among 2𝒥 parent-offspring pairs in the whole sample.

#### A.6 Proof of Theorem 3

##### Part A

We first discuss the covariance between siblings within the same family. Let 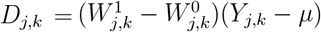 for offspring *k* in family *j* and 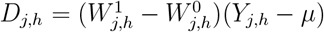 for offspring *h* in family *j*. Here we derive a closed formula for ℂ (*D*_*j,k*_, *D*_*j,h*_|*N*) where *k* ≠ *h*. We show that when the null hypothesis is true ℂ (*D*_*j,k*_, *D*_*j,h*_|*N*) = 0 and in general ℂ (*D*_*j,k*_, *D*_*j,h*_|*N*) ≥ 0. Let 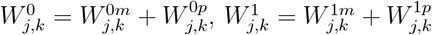 where

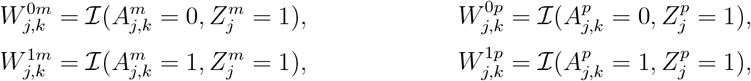

and also 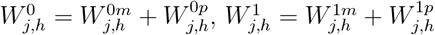 where

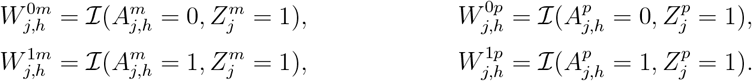

Then

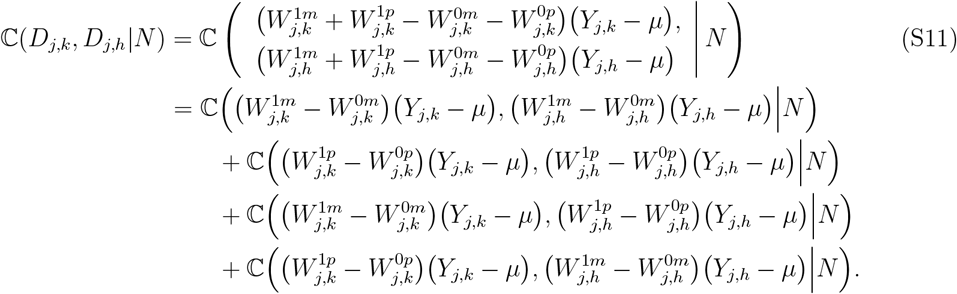

Eq. (S11) involves four covariances whose calculations follow the same algebra. For the first covariance,

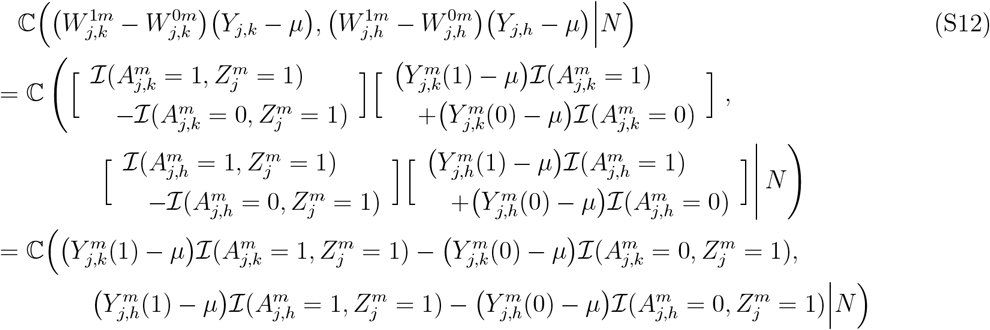

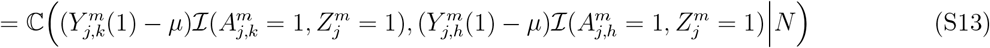

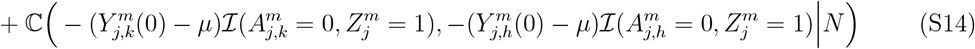

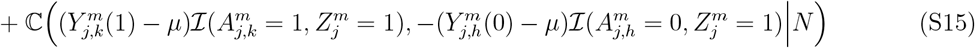

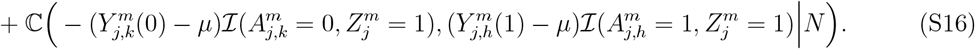

We first calculate Eq. (S13) as follows.

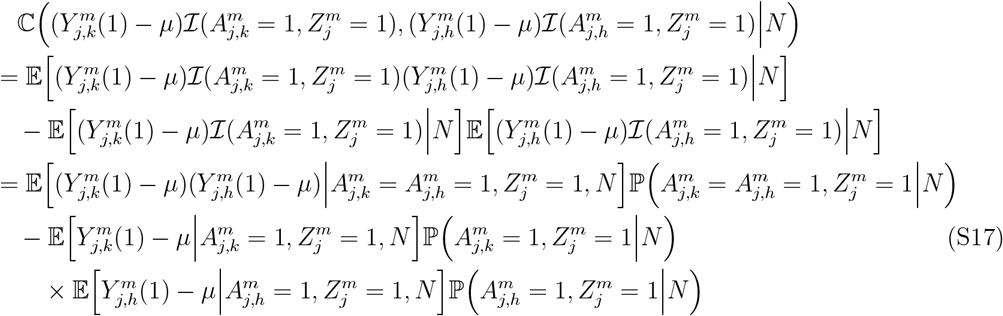

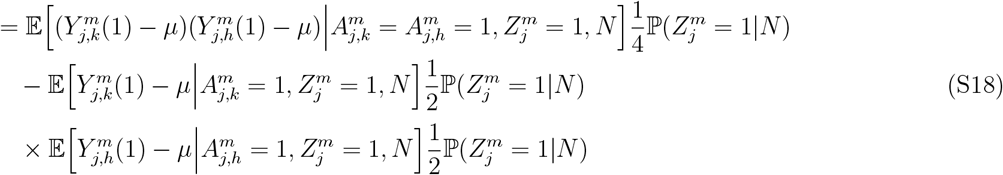

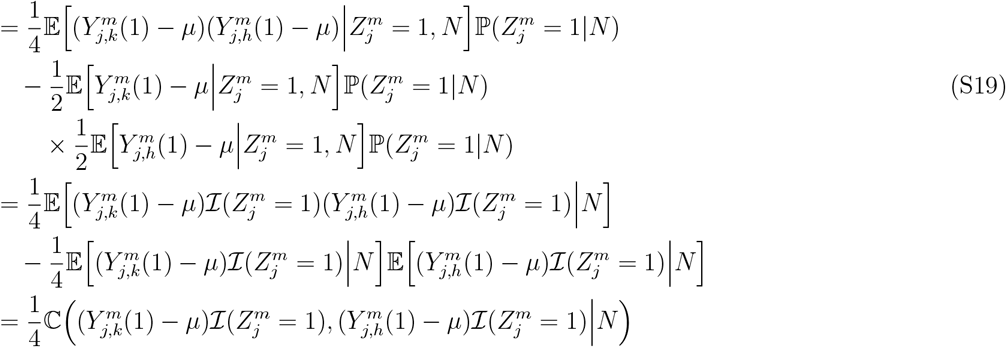

Eq. (S17) to Eq. (S18) follows from

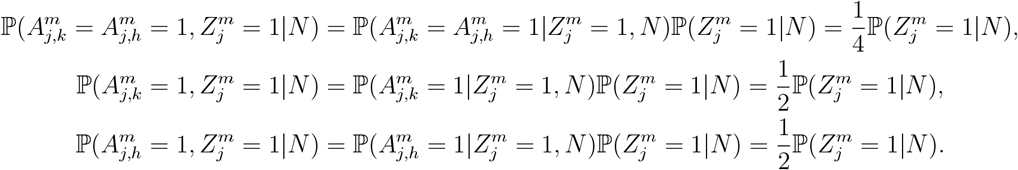

Eq. (S18) to Eq. (S19) follows from Lemma 1 that 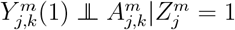 and 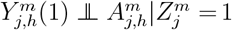. We use the same approach to calculate the other three covariances in Eq. (S12) as follows.

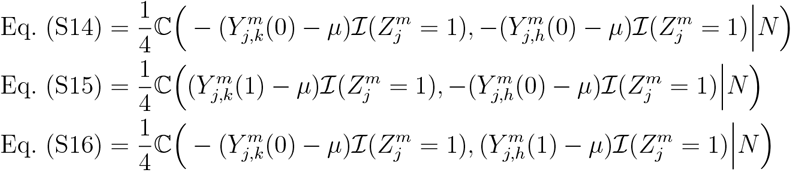

We plug these four covariances into Eq. (S12) to derive the following covariance.

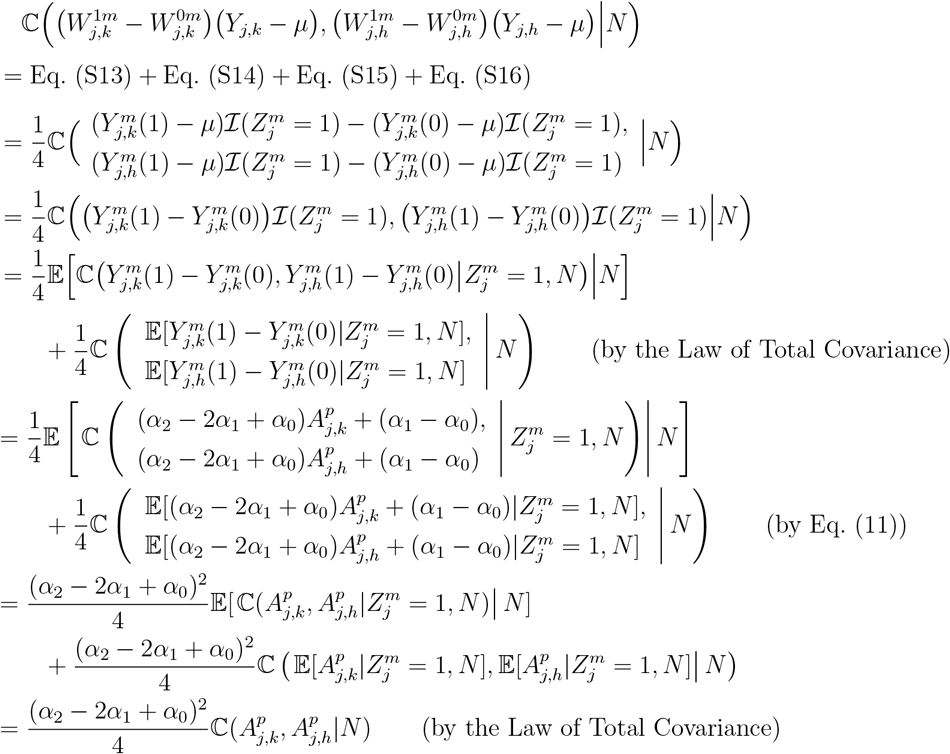

We use the same approach to calculate the other three covariances in Eq. (S11) and add these four covariances to derive ℂ (*D*_*j,k*_, *D*_*j,h*_|*N*) as follows.

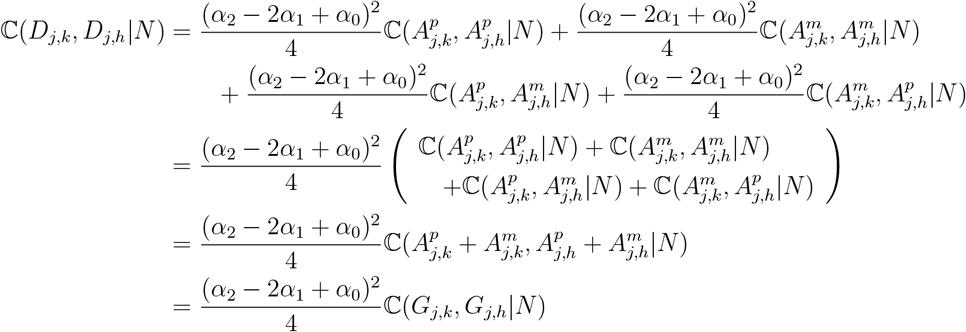

When the null hypothesis of no causality is true, *α*_0_ = *α*_1_ = *α*_2_. Therefore, (*α*_2_ − 2*α*_1_ + *α*_0_) = 0 and ℂ (*D*_*j,k*_, *D*_*j,h*_|*N*) = 0 when the null hypothesis of no causality is true. In general, (*α*_2_ − 2*α*_1_ + *α*_0_)^2^ ≥ 0. The property that ℂ (*G*_*j,k*_, *G*_*j,h*_|*N*) ≥ 0 is satisfied in prevalent population genetics models including the Identical-By-Descent (IBD) model, the co-ancestry model, etc [29]. In this case, ℂ (*D*_*j,k*_, *D*_*j,h*_|*N*) ≥ 0 in general.

##### Part B

We now consider the covariance between offspring from different families. Let 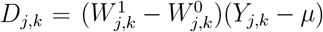 for offspring *k* in family *j* and 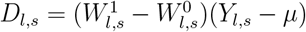 for offspring *s* in family *l*. Here we derive a closed formula for ℂ (*D*_*j,k*_, *D*_*l,s*_|*N*) where *j*≠ *l*. We show that when the null hypothesis is true, ℂ (*D*_*j,k*_, *D*_*l,s*_|*N*) = 0 and in general ℂ (*D*_*j,k*_, *D*_*l,s*_|*N*) ≥ 0. For offspring *k* in family *j*, let 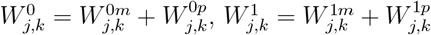 where

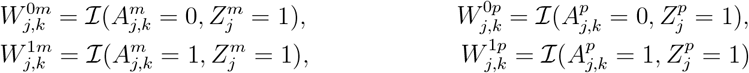

For offspring *s* in family *l*, let 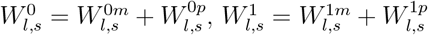 where

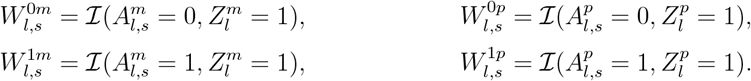

Then

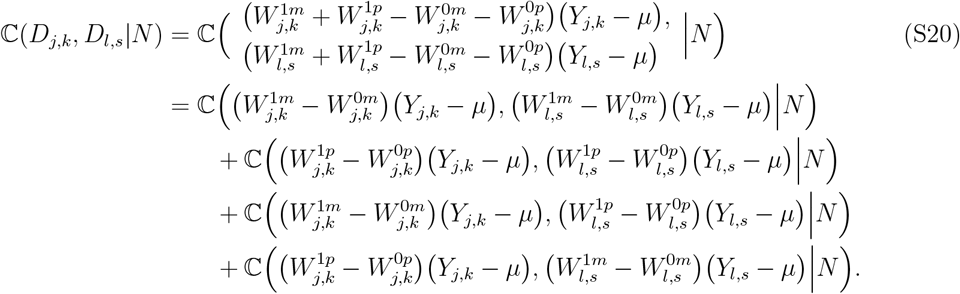

Eq. (S20) involves four covariances whose calculations follow the same algebra. We calculate the first covariance as follows.

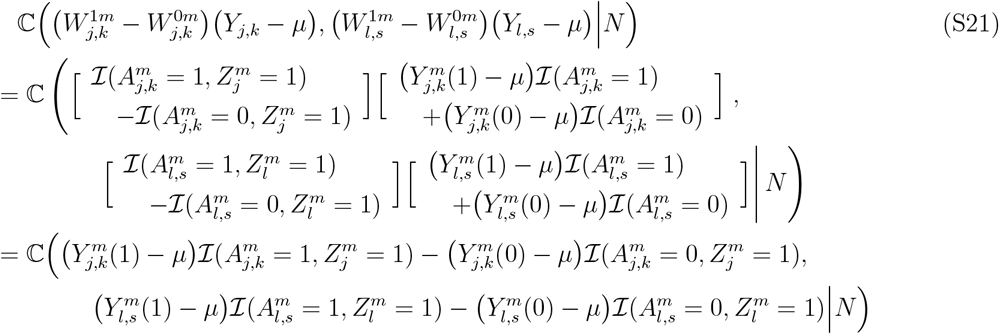

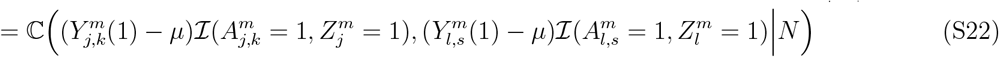

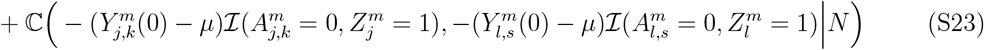

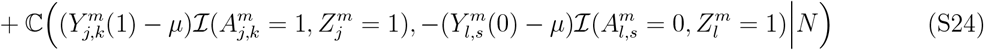

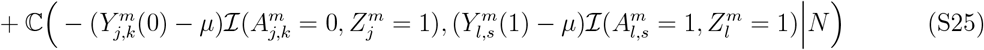

We first calculate Eq. (S22) as follows.

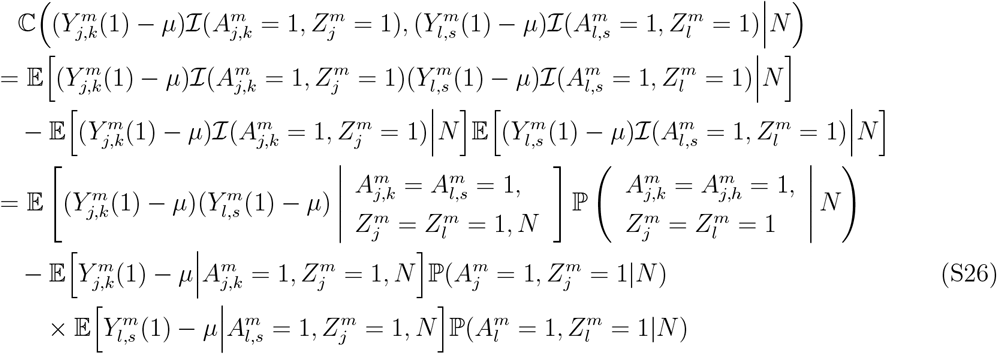

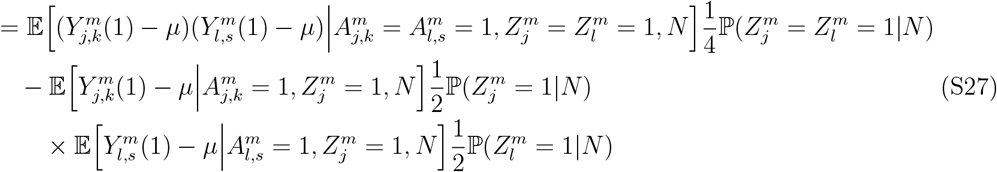

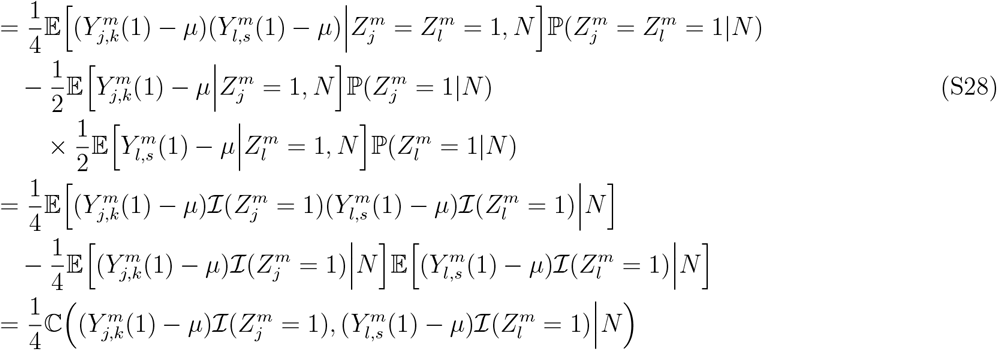

From Eq. (S26) to Eq. (S27) follows that

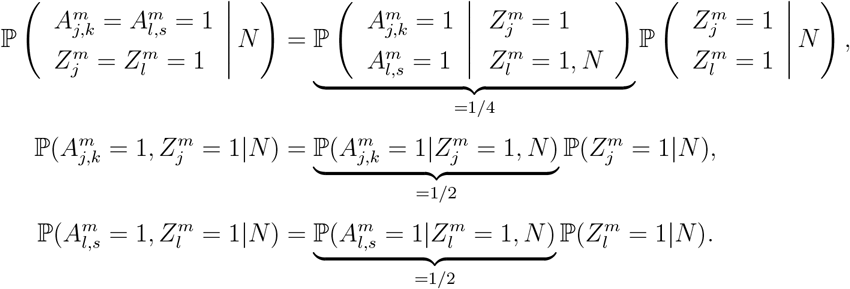

Eq. (S27) to Eq. (S28) follows from Lemma 1 that 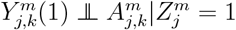 and 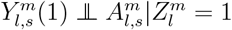. We use the same approach to calculate the other three covariances in Eq. (S12) as follows.

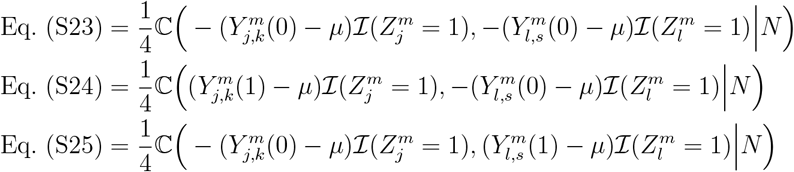

We plug these four covariances into Eq. (S21) to derive the following.

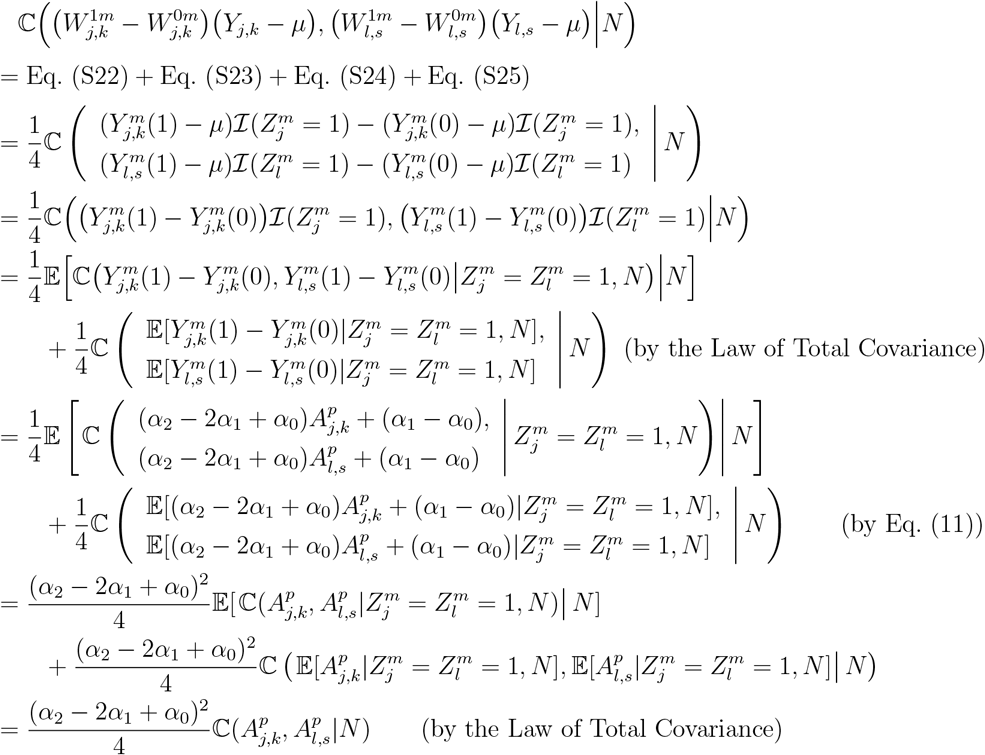

We use the same approach to calculate the other three covariances in Eq. (S20) and add these four covariances to derive ℂ (*D*_*j,k*_, *D*_*l,s*_|*N*) as follows.

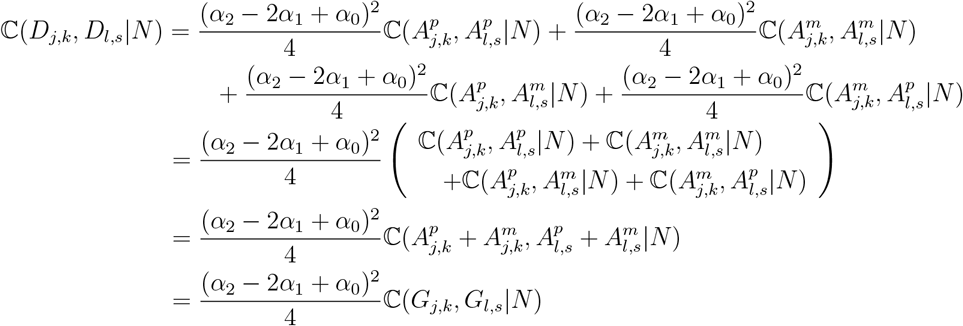

When the null hypothesis of no causality is true, *α*_0_ = *α*_1_ = *α*_2_. Therefore, (*α*_2_ − 2*α*_1_ + *α*_0_) = 0 and ℂ (*D*_*j,k*_, *D*_*l,s*_|*N*) = 0 when the null hypothesis of no causality is true. In general, (*α*_2_−2*α*_1_+*α*_0_)^2^ ≥ 0. Besides, ℂ (*G*_*j,k*_, *G*_*l,s*_|*N*) ≥ 0 is assumed to be satisfied in prevalent population genetics models including the IBD model, the co-ancestry model, etc. Then ℂ (*D*_*j,k*_, *D*_*l,s*_|*N*) ≥ 0 in general.

##### Part C

We derive a closed formula for 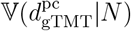 and use Part A and B to complete the proof of Theorem 3. Let 𝒥 be the total number of offspring in that 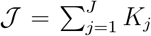. Recall that 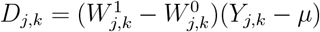 for offspring *k* in family *j*. Then 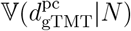 equals

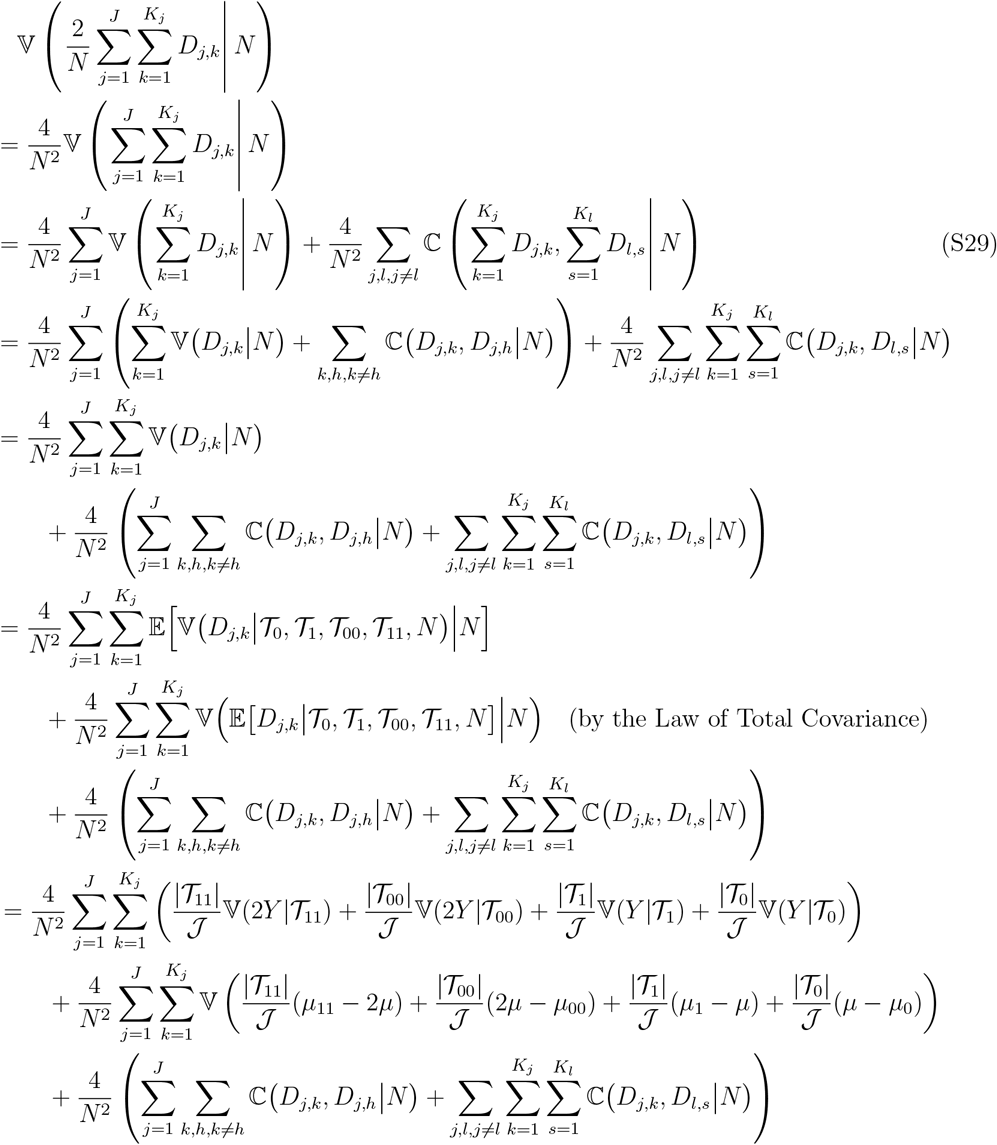

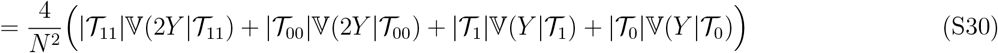

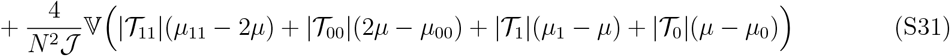

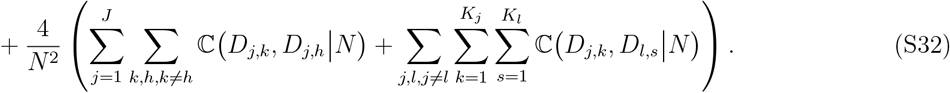

We have the following key observations:

A. For Eq. (S31), when the null hypothesis of no causality is true, *µ*_11_ = *µ*_00_ = 2*µ* and *µ*_1_ = *µ*_0_ = *µ* so that (S31) equals zero. When the alternative hypothesis is true, (S31) is a variance so it is non-negative.
B. For Eq. (S32), by Part A and B in Appendix A.6, (S32)= 0 when the null hypothesis of no causality is true, it follows that (S32)≥ 0 when the alternative hypothesis is true.

Therefore, when the null hypothesis that *δ*_TMT_ = 0 is true, 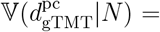 Eq. (S30). In general, 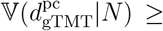 Eq. (S30). Notice that Eq. (S30) is exactly the right hand side of Eq. (14) in Theorem 3. This completes the proof of Theorem 3.

### B Simulations

#### B.1 Simulating genotypes

Both parental genotypes matrices ***Z***^*m*^ and ***Z***^*p*^ were sampled from a structured population based on a standard admixture model [27, 28] with *K* = 4 admixed populations. We configured this population to have *F*_ST_ = 0.2 by utilizing the bnpsd R package [29, 40]. We simulated the ancestral allele frequencies from the Uniform(0.1, 0.9), which is an option in the bnpsd R package.

#### B.2 Simulating quantitative traits

We first generated a non-genetic factor associated with the population structure. Let ***E*** = {*E*_*j*_} be the random non-genetic factor. We adopted the admixture proportion ***q*** = {*q*_*ju*_} from Appendix B.1 to simulate *E*_*j*_ as

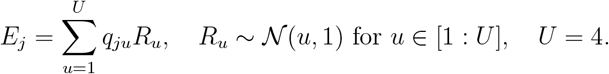

Let *C* be the set of causal SNPs. We followed Eq. (9) to simulate the child’s trait ***Y*** = {*Y*_*j,k*_} as

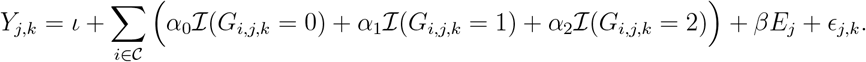

To show that the above equation satisfies the trait model in Eq. (9), at causal SNP *c* (*c* ∈ 𝒞), rewrite the equation as

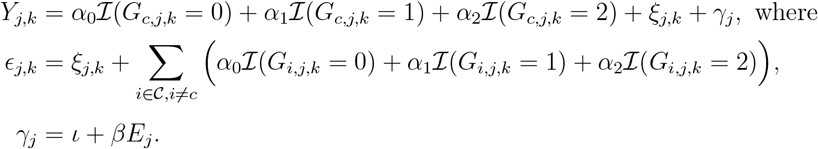

Note that the first line satisfies Eq. (9). We considered the following parameters

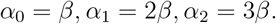

Then the ratio (*α*_2_ − *α*_1_)*/*(*α*_1_ − *α*_0_) is 1. Given the desired heritability 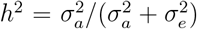 where we set 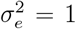, we simulated *β* such that 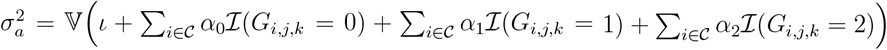, achieving the desired *h*^2^.

#### B.3 Simulating family-specific confounding effects

We followed Appendix B.1 to simulate genotypes of 1,500 nuclear families (500 trios, 500 tetrads and 500 quintets). This leads to 3,000 offspring and 3,000 parents with 100,000 SNPs per individual. We randomly chose 100 causal SNPs and denoted them as the causal SNP set 𝒞. We randomly chose a target causal locus *i*^∗^ from 𝒞 for testing. Let ℐ (·) be the indicator function. We generated a family-specific confounding effect 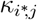 that is correlated with the parental homozygous genotypes at locus *i*^∗^ by 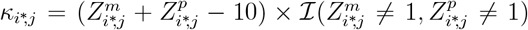. We generated child phenotypes according to

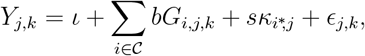

where we generated *ϵ*_*j,k*_ from Normal 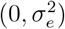 and we set 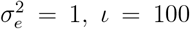 . The coefficient *b* is determined such that 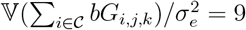.

### C UK Biobank Data Analysis

#### C.1 Identifying trios

We followed the criteria in `KING` [31] to identify 28,895 parent-child pair candidates that have estimated kinship around 1/4 (between 2^−5*/*2^ and 2^−3*/*2^) and a discordant homozygote rate less than 0.1. We filtered pairs whose age difference is less than 17, leaving 4,494 mother-child pairs and 1,998 father-child pairs. We excluded four mother-child pairs that share the same child because the mothers are twins. Among the remaining parent-child pairs, we detected nuclear families that have both parents available and at least one child, leading to 990 trios (two parents and one child) and 37 quartets (two parents and two offspring) with 1,064 distinct offspring and 2,054 parents in total.

#### C.2 Blood pressure phenotypes quality control

We used two original phenotypes in UK Biobank including systolic blood pressure (SBP, field id p4080) and diastolic blood pressure (DBP, field id p4079). These are automated reading results and each phenotype contains two repeated measures per individual that were taken a few moments apart. We calculated the mean of two repeated measures per individual as the target trait. When automated reading results were missing, we used manual reading measures of systolic blood pressure (field id p93) and diastolic blood pressure (field id p94) to fill missing values. Among our study cohort, 1,062 offspring have valid blood pressure records and two offspring have neither automated nor manual reading results. We followed a previous QC pipeline [41] to adjust SBP and DBP for the use of blood pressure lowering medication.

### D Supplementary Figures and Tables

**Figure S1:**
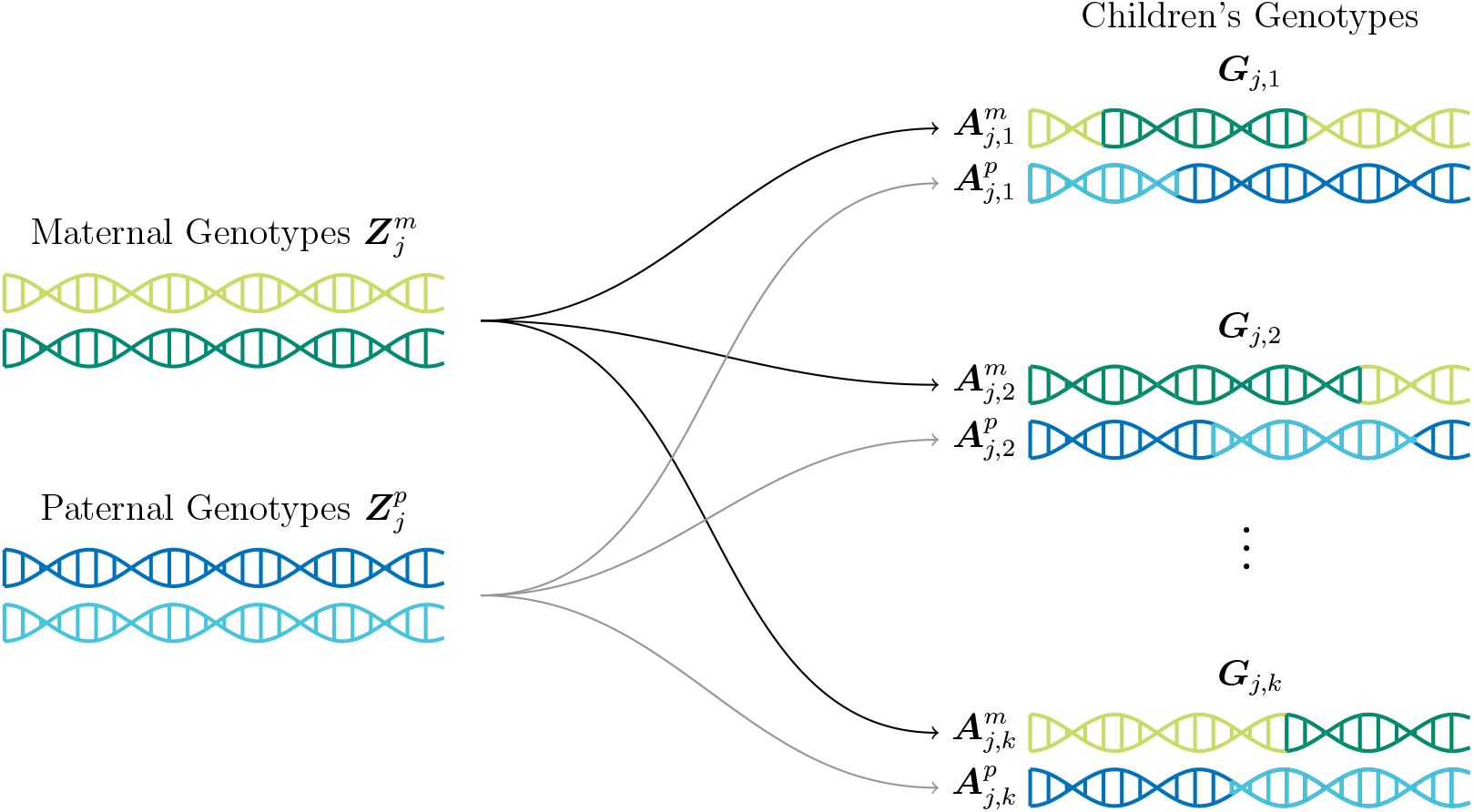
Schematic of randomized genetic transmission in a nuclear family.

**Table S1:** The conditional probability of observing an offspring genotype given parental genotypes.

| Offspring Genotype<br>$G$ | Parental Genotypes<br>$(Z^m, Z^p)$ | Conditional Probability<br>$\mathbb{P}(G\vert Z^m, Z^p)$ |
| --- | --- | --- |
| 2 | (2,2) | 1 |
| 2 | (2,1) | 1/2 |
| 2 | (1,2) | 1/2 |
| 2 | (1,1) | 1/4 |
| 1 | (2,1) | 1/2 |
| 1 | (1,2) | 1/2 |
| 1 | (2,0) | 1 |
| 1 | (0,2) | 1 |
| 1 | (1,1) | 1/2 |
| 1 | (1,0) | 1/2 |
| 1 | (0,1) | 1/2 |
| 0 | (1,1) | 1/4 |
| 0 | (1,0) | 1/2 |
| 0 | (0,1) | 1/2 |
| 0 | (0,0) | 1 |

**Table S2:** The conditional probability of observing two offspring genotypes given parental genotypes.

| Sibling Pairs' Genotypes<br>$(G_{j,k}, G_{j,h})$ | Parental Genotypes<br>$(Z_j^m, Z_j^p)$ | Conditional Probability<br>$\mathbb{P}(G_{j,k}, G_{j,h} \vert Z_j^m, Z_j^p)$ |
| --- | --- | --- |
| (2,2) | (2,2) | 1 |
| (2,2) | (2,1) | 1/4 |
| (2,2) | (1,2) | 1/4 |
| (2,2) | (1,1) | 1/16 |
| (2,1) | (2,1) | 1/4 |
| (2,1) | (1,2) | 1/4 |
| (2,1) | (1,1) | 1/8 |
| (1,2) | (2,1) | 1/4 |
| (1,2) | (1,2) | 1/4 |
| (1,2) | (1,1) | 1/8 |
| (1,1) | (2,1) | 1/4 |
| (1,1) | (1,2) | 1/4 |
| (1,1) | (2,0) | 1 |
| (1,0) | (1,1) | 1/8 |
| (1,0) | (1,0) | 1/4 |
| (1,0) | (0,1) | 1/4 |
| (0,1) | (1,1) | 1/8 |
| (0,1) | (1,0) | 1/4 |
| (0,1) | (0,1) | 1/4 |
| (0,0) | (1,1) | 1/16 |
| (0,0) | (1,0) | 1/4 |
| (0,0) | (0,1) | 1/4 |
| (0,0) | (0,0) | 1 |

